# Transcriptomics, metabolomics and lipidomics of chronically injured alveolar epithelial cells reveals similar features of IPF lung epithelium

**DOI:** 10.1101/2020.05.08.084459

**Authors:** Willy Roque, Karina Cuevas-Mora, Dominic Sales, Wei Vivian Li, Ivan O. Rosas, Freddy Romero

**Affiliations:** Department of Medicine, Rutgers – New Jersey Medical School, 185 S Orange Ave, Newark, NJ 07103, USA; Department of Medicine, Division of Pulmonary, Allergy and Critical Care and the Center for Translational Medicine; the Jane & Leonard Korman Respiratory Institute, Philadelphia, PA, US; Department of Biostatistics and Epidemiology, Rutgers School of Public Health, Piscataway 08854; Pulmonary, Critical Care and Sleep Medicine, Baylor College of Medicine

**Keywords:** mitochondrial dysfunction, heat shock proteins, proteostasis, cellular senescence, idiopathic pulmonary fibrosis, metabolism

## Abstract

The current hypothesis suggests that Idiopathic pulmonary fibrosis (IPF) arises as a result of chronic injury to alveolar epithelial cells and aberrant activation of multiple signaling pathways. Dysfunctional IPF lung epithelium manifests many hallmarks of aging tissues, including cellular senescence, mitochondrial dysfunction, metabolic dysregulation, and loss of proteostasis. Unfortunately, this disease is often fatal within 3-5 years from diagnosis, and there is no effective treatment. One of the major limitations to the development of novel treatments in IPF is that current models of the disease fail to resemble several features seen in elderly IPF patients. In this study, we sought to develop an *in vitro* epithelial injury model using repeated low levels of bleomycin to mimic the phenotypic and functional characteristics of the IPF lung epithelium. Consistent with the hallmarks of the aging lung epithelium, we found that chronic-injured epithelial cells exhibited features of senescence cells, including an increase in β-galactosidase staining, induction of p53 and p21, mitochondrial dysfunction, excessive ROS production, and proteostasis alteration. Next, combined RNA sequencing, untargeted metabolomics, and lipidomics were performed to investigate the dynamic transcriptional, metabolic, and lipidomic profiling of our *in vitro* model. We identified that a total of 8,484 genes with different expression variations between the exposed group and the control group. According to our GO enrichment analysis, the down-regulated genes are involved in multiple biosynthetic and metabolic processes. In contrast, the up-regulated genes in our treated cells are responsible for epithelial cell migration and regulation of epithelial proliferation. Furthermore, metabolomics and lipidomics data revealed that overrepresented pathways were amino acid, fatty acid, and glycosphingolipid metabolism. This result suggests that by using our *in vitro* model, we were able to mimic the transcriptomic and metabolic alterations of those seen in the lung epithelium of IPF patients. We believe this model will be ideally suited for use in uncovering novel insights into the gene expression and molecular pathways of the IPF lung epithelium and performing screening of pharmaceutical compounds.

## Introduction

Idiopathic pulmonary fibrosis (IPF) represents one of the most aggressive and irreversible lung diseases, usually diagnosed in the fifth decade of life, carries a very poor prognosis, an unknown etiology, and limited therapeutic options (9, 23, 36). IPF is characterized by chronic alveolar epithelial injury, leading to the development of complex pro-fibrotic cascade activation and ending with extracellular matrix deposition. The development of an effective treatment for pulmonary fibrosis will require an improved understanding of its molecular pathogenesis (4, 45).

Multiple risk factors have been related to the IPF pathogenesis, including but not limited to advanced age, genetic factors, and environmental, such as cigarette smoke factors and dust. Dysfunctional IPF lung epithelium manifests many hallmarks of aging tissues, including cellular senescence, mitochondrial dysfunction, loss of proteostasis, and metabolic dysregulation (2, 22, 41, 42, 55).

Alveolar epithelial type II (AE2) cells have been shown to undergo senescence in IPF lungs and could contribute to the activation of pulmonary fibroblast via increasing the expression of senescence-associated secretory phenotype (SASP) (7, 17, 21, 53). Moreover, recent studies have shown that age-dependent accumulation of senescent cells may block fibrosis resolution following bleomycin exposure and that targeting senescent cells for destruction can be quite effective in ameliorating experimentally induced respiratory conditions, highlighting a potential role for anti-senescent approaches in the treatment of chronic lung disease (13, 15, 19).

Emerging evidence indicates that mitochondrial dysfunction and metabolic reprogramming occur in a wide range of respiratory disorders, including a variety of fibrotic lung conditions (15, 29, 39). For example, we recently demonstrated that diverse types of profibrotic insults (bleomycin, radiation, silica) induce similar metabolic changes in the alveolar epithelium of the lung (51). These changes include decreased ATP production and a shift toward the use of glycolysis as a means for driving cellular metabolism. Similarly, proteostasis collapse has also been described in the lung of patients with IPF, indicating that dysregulation in protein homeostasis is also a feature of human fibrotic lung conditions (38, 40, 43).

Although it is recognized that epithelial dysfunction underlies the development of the disease, there is limited information regarding the mechanism by which typical environmental levels of exposure can contribute to the onset of this disease. Understanding the causes and elucidate the mechanisms associated with the pathogenesis of the disease thereof has been limited by the lack of an *in vitro* model. Existing *in vitro* systems used to model IPF rely on acute exposures to agents such as bleomycin, silica, or asbestos dust (5, 52). However, these models are defective due to the fact that they fail to mimic the chronic, repetitive insults seen by the IPF lung epithelium. In this study, our objective was the development of a novel *in vitro* model that better resembles the IPF lung epithelium. Our investigation includes a transcriptomic analysis allowing identification of gene networks involved in response to repeated low-level bleomycin exposure. Also, we combined metabolomics and lipidomic analysis to identify the metabolic and lipidomic sub-network involved in the biological response to low levels of bleomycin exposure. Our study is the first in-depth investigation of the molecular and biochemistry profiles underlying the effects of repeated low-level bleomycin exposure of AE2 cells, and by integrating the transcriptome, metabolome, and lipidomic, we showed striking similarities with the IPF lung epithelium. We believe this model will be useful in understanding the early biological mediators of lung fibrosis and as a model to evaluate pharmaceutical compounds.

## Methods

### Cell culture and reagents

Mouse MLE12 cells were obtained from the American Type Culture Collection (ATCC, CRL-2110). We created an innovative *in vitro* model of chronic bleomycin exposure using murine alveolar type-II epithelial cells. On day 0, MLE-12 cells were treated with 10 μ/ml bleomycin (Enzo life science, BML-AP302-0050) for 24h. Day 2, bleomycin was removed, and the culture medium was refreshed. Day 4, cells were treated again with 10 μg/ml bleomycin. Day 5, bleomycin was subsequently removed, and cells were collected at day 7. Cellular senescence markers were evaluated on day seven after the initiation of bleomycin injury.

### Mitochondrial ROS assessments

Mitochondrial ROS was detected by incubating cells with 5 μM Mito-Sox Red (Invitrogen, Carlsbad, CA) for 10 min at 37 C and measuring fluorescence using a microplate reader.

### Mitochondrial DNA damage assay

Nuclear and mitochondrial (mt) DNA damage was assessed by qPCR as previously described. In brief, genomic DNA was isolated using the Qiagen Genomic-Tip 20G and Qiagen DNA Buffer kit. PCR was performed to amplify both short and long forms of mtDNA fragment and nuclear DNA (β-globin). Each DNA was quantified using Pico-Green (Thermo Fisher Scientific, Waltham, MA) and a microplate fluorescent reader (PerkinElmer, Waltham, MA.) at excitation and emission wavelengths of 485 and 530 nm, respectively. Data obtained from the mitochondrial small fragment was used to normalize the results of the mitochondrial long fragment. The numbers of mitochondrial lesions were calculated by using the following equation: D= (1-2 (−Δlong−Δshort) × 10,000 (bp)/size of the long fragment (bp).

### Oxygen consumption measurements

Oxygen consumption rate (OCR) was measured using the Seahorse XFp Bioanalyzer (Seahorse Bioscience). In brief, mouse MLE12 cells were seeded at a concentration of 20,000 cells/well on XFp cell plates (Seahorse Bioscience, 103057-100) 24 hours prior to the initiation of studies. The concentration of FCCP (also known as trifluoromethoxy carbonylcyanide phenylhydrazone), antimycin A, and rotenone were 0.75μM, 2μM, and 2μM respectively. Pyruvate (10 mM) and glucose (25 mM) served as a substrate. The final concentrations of injected compounds were as follows: port A, 2uM oligomycin; port B 0.75 FCCP, port C 2uM antimycin and 2uM rotenone.The protocol and algorithm for our XF-PMP assay were designed using wave 2.4 software.

### Senescence-associated beta-galactosidase (SA-β-gal) detection

SA-β-gal staining was performed using the β-galactosidase kit (Dojindo Molecular Technology, 1824699-57-1) according to the manufacturer’s instructions. To prevent false positives, experiments were performed on cells at 70% confluence.

### Protein aggregation assay

A Proteostat aggresome detection kit (Enzo Life Sciences, ENZ-51035-0025) was used to characterize the aggregates and aggresomes in MLE12 cells. All components of this kit were prepared according to the manufacturer’s instructions. Monolayer cells were grown on cell culture dishes. Cells were washed with 1X PBS and fixed with 4% paraformaldehyde for 10 min at room temperature. After washed with excess 1 X PBS, cells were permeabilized with 0.5% Triton X-100 (Millipore-Sigma, 9002-931), 3 mM EDTA (Sigma-Aldrich, 03690) in 1 X PBS for 30 min. Cells were again washed twice with 1X PBS and the ProteosStat dye was added at 1:2000 dilution for 30 min.

### Proteasome activity

MLE12 cells were washed twice with cold PBS, collected with a rubber policeman, and lysed in ice-cold proteasome buffer (10 mM Tris, pH 7.8, 1 mM EDTA, 5 mM MgCl_2_, 5 mM ATP). Cell debris was removed by centrifugation for 15 min at 14,000 x g, and the supernatant was used for the assay. Chymotrypsin-like, trypsin-like and caspase-like proteasome activity was assayed at 37 °C using 10 μg of protein extracts in proteasome buffer in the presence of 100 μM Suc-LLVY-aminoluciferin (Succinyl-leucine-valine-tyrosine-aminoluciferin), Z-LRR-aminoluciferin (Z-leucine-arginine-arginine-aminoluciferin and Z-nLPnLD-aminoluciferin(Z-norleucine-proline-norleucine-aspartate-aminoluciferin), respectively (Promega Corporation, G8531). The reaction was monitored by fluorimetric measurements every 10 min for 1h (excitation 350 nm, emission 460 nm) in a Synergy HT Multi-Detection microplate reader. Proteasome activity was determined as the difference between the total activity of cell lysis and the remaining activity in the presence of 20 μM MG132 (Millipore-Sigma, M7449).

### Western blot analysis

Controls and bleomycin-injured MLE12 cells were lysed in ice-cold buffer (PBS, 0.05% Tween 20, pH 7.4) containing protease inhibitors (Active motif, 37491) and phosphatase inhibitor (Active motif, 37493). After centrifugation (14,000 x g, 10 min, 4°C), the supernatant was collected. Fibroblast cell nuclear fraction was extracted using a commercially available kit (Active motif, 40010) according to the manufacturer’s instructions. Protein concentration was determined by Pierce™ BCA assay kit (ThermoFisher, 23225). Protein samples (20 μg) were solubilized in 4 × Laemmli sample buffer, heated at 95°C for 10 min, centrifuged at 3,000 g for 1 min, loaded on a 10% Tris-HCl-SDS-polyacrylamide gel and run for 1 h at 100 V. Protein was transferred to a nitrocellulose membrane (ThermoFisher, LC2000) and then blocked with Odyssey Blocking Buffer (Li-Cor Biosciences, 927-50000) for 1 h at room temperature followed by incubation overnight at 4°C with a specific primary antibody to p-p53^ser15^ (Cell Signaling, 9284S), β-actin (Cell Signaling, 84575S), Lamin B1(Cell Signaling, 13435S), γ-H2AX (Cell Signaling, 2577S), HSP90 (Cell Signaling, 4877S), HSP70 (Cell Signaling, 4873S), HSP40 (Cell Signaling, 48715), HSF1(Cell Signaling, 12972S), p-HSF1^ser307^ (Abcam, ab47369, HSF^K298 sumo^ (MyBioSource, MBS9211638), p21(Cell Signaling, 2946S), at a dilution of 1:1,000 in blocking buffer with 0.1% Tween-20, followed by incubation with Donkey anti-Rabbit (Li-Cor Biosciences, 926-32213) or anti-mouse secondary antibody (Li-Cor Biosciences,926-32212) at a dilution of 1:8,000 in blocking buffer. After three washings with PBS, immunoblots were visualized using the Odyssey infrared imaging system (Li-Cor Biosciences).

### RNA isolation and Real-time quantitative PCR (qRT-PCR)

Gene transcript levels were quantified by real-time PCR. Total RNA was isolated using RNeasy Mini-Kit (QIAGEN, 74104), according to the manufacturer’s instructions. A summary of primer sets manufactured by Integrated DNA Technologies can be found in supplemental Table 1. Relative gene expression levels were quantified with the Livak method (2-ΔΔCT) by normalizing expression of the gene of interest to Hprt, demonstrated to be unaffected by bleomycin, and calculating the relative fold differences between experimental and control samples. Assays were completed in technical triplicate.

### RNA sequence library construction

Five million cells were used to extract total RNA using the RNeasy Mini Kit (Qiagen) according to the manufacturer’s protocol. After quantification of RNA by Fragment Analyzer (Advanced Analytical), 1.5 μg of total RNA was used to construct sequencing libraries by TruSeq RNA Sample Preparation Kit (Illumina) according to the manufacturer’s instructions

### RNA sequence analysis

RNA-seq was performed with Illumina Hiseq 2000 by Novogene, USA. Downstream analysis was performed using a combination of programs including STAR, HTseq, Cufflink and our wrapped scripts. Alignments were parsed using Tophat program, and differential expressions were determined through DESeq2/edgeR. GO and KEGG enrichment were implemented by the ClusterProfiler. Gene fusion and difference of alternative splicing events were detected by Star-fusion and rMATS software. For DESeq2 with biological replicates). Differential expression analysis between two conditions/groups (two biological replicates per condition) was performed using the DESeq2 R package (2_1.6.3). DESeq2 provides statistical routines for determining differential expression in digital gene expression data using a model based on the negative binomial distribution. The resulting P-values were adjusted using the Benjamini and Hochberg’s approach for controlling the False Discovery Rate (FDR). Genes with an adjusted P-value <0.05 found by DESeq2 were assigned as differentially expressed. For edgeR without biological replicates) prior to differential gene expression analysis, for each sequenced library, the read counts were adjusted by edgeR program package through one scaling normalized factor. Differential expression analysis of two conditions was performed using the edgeR R package (3.16.5). The P values were adjusted using the Benjamini & Hochberg method. Corrected P-value of 0.05 and absolute fold change of 1 was set as the threshold for significantly differential expression. The Venn diagrams were prepared using the function Venn diagram in R based on the gene list for a different group.

Next, we conducted a principal component analysis (PCA) to compare the overall transcriptomic profiles in our data to those from Xu et al. (56) and Nance et al. (32). The PCA approach was applied to the FPKM (fragments per kilobase of transcript per million mapped reads) values of gene expression after log2 transformation. To reduce the batch effect in samples sequenced in different studies, we applied the limma method on the gene expression data before applying PCA (49).To further identify the genes that are jointly differentially expressed (DE) between the control samples and the treated/IPF samples, we also conducted a differential expression analysis using the DESeq2 method (25). To study the biological functions of the DE genes, we performed the Gene Ontology (GO) enrichment analysis in the 100 most significant up-regulated and down-regulated genes, respectively (58).

### Metabolomics analysis

Samples were thawed before applied to extraction procedure and extracted with 800 μl of 80% methanol, 5 μl of DL-o-Chlorophenylalanine (2.8 mg/mL) and 5 μl of LPC (12:0) with ultrasound for 30 min. Then all samples were kept at −20 °C for 1 h. After that, samples were vortex for 30 s, and centrifuged at 12000 rpm and 4 °C for 15 min. Finally, 200 μl of supernatant was transferred to a vial for LC-MS analysis by Creative Proteomics, Inc. Quality control (QC) samples were used to evaluate the methodology. The same amount of extract was obtained from each sample and mixed as QC samples. The QC sample was prepared using the same preparation procedure. There are 1 pair of analyses in total between the control group and the senescent group. The separation was performed by Ultimate 3000LC combined with Q Exactive MS (Thermo) and screened with ESI-MS (targeted MS/MS mode). The LC system is comprised of a Thermo Hyper gold C18 (100×2.1mm 1.9 μm) with Ultimate 3000LC. The mobile phase is compose of solvent A (0.1% formic acid-5% acetonitrile-water) and solvent B (0.1% formic acid-acetonitrile) with a gradient elution (0-1.5 min, 100-80% A; 1.5-9.5 min, 80-0% A; 9.5-14.5 min, 0% A; 14.5-14.6 min, 0-100% A; 14.6-18 min, 100% A). The flow rate of the mobile phase is 0.35 mL·min-1. The column temperature is maintained at 40°C, and the sample manager temperature is set at 4°C.

Mass spectrometry parameters in ESI+ and ESI− mode are listed as follows: ESI+: Heater Temp 300 °C; Sheath Gas Flow rate, 45 arb; Aux Gas Flow Rate, 15 arb; Sweep Gas Flow Rate, 1 arb; spray voltage, 3.0 kV; Capillary Temp, 350 °C; S-Lens RF Level, 30%. ESI−: Heater Temp 300 °C, Sheath Gas Flow rate, 45 arb; Aux Gas Flow Rate, 15arb; Sweep Gas Flow Rate, 1 arb; spray voltage, 3.2 kV; Capillary Temp,350 °C; S-Lens RF Level, 60%. At the beginning of the sequence, we run four quality control (QC) samples to avoid small changes in both chromatographic retention time and signal intensity. The QC samples are also injected at regular intervals (every ten samples) throughout the analytical run.

The raw data are acquired and aligned using the Compound Discover (3.0, Thermo) based on the m/z value and the retention time of the ion signals. Ions from both ESI− and ESI+ are merge and import into the SIMCA-P program (version 14.1) for multivariate analysis. A Principal Components Analysis (PCA) is first used as an unsupervised method for data visualization and outlier identification. Supervised regression modeling is then performed on the data set by use of Partial Least Squares Discriminant Analysis (PLS-DA) or Orthogonal Partial Least SquaresDiscriminant Analysis (OPLS-DA) to identify the potential biomarkers. The biomarkers are filtered and confirmed by combining the results of the VIP values (VIP>1.5) and t-test (p < 0.05). The quality of the fitting model can be explained by R2 and Q2 values. R2 displays the variance explained in the model and indicated the quality of the fit. Q2 displays the variance in the data, indicating the model’s predictability.

To compare the metabolomics profiles of our in vitro model with those of the IPF patients, we collected the metabolomics data from an IPF study with 15 controls and 13 IPF patients. The data available from Kang et al. only contains summary statistics of the metabolites but not individual observations, so we used the following approach for the comparison (18). First, we restrict our comparison to the 23 metabolites that were shown to have significant changes (p-value < 0.01) between control and IPF samples. For these metabolites, we first summarized their intensity in our samples and then scaled the intensity to zero mean and unit variance. Next, we used the two-sample t-test to compare the scaled intensity of these metabolites between the control and treated cells. Finally, we compared which metabolites demonstrate consistently significant changes in our and Kang et al.’s data.

### Untargeted lipidomics

Chemical extraction was carried out on 12 samples (six biological replicates for each experimental group. Samples were extracted with 1.5mL Chloroform:MeOH (2:1,v/v) and 0.5mL ultrapure water and vortex for 1min. Samples were centrifuge 10 min at 3,000 rpm 4°C, and the lower phase was transferred to a new tube for evaporation. The dried extract was reconstituted with 400 μl of isopropyl alcohol:MeOH (1:1,v/v) and centrifuge for 10 min at 12,000 rpm, 4°C. Finally, the resulting supernatant was analyzed using a LC-MS/MS system by Creative Proteomics, Inc. The separation was performed by UPLC (Thermo, Ultimate 3000LC) and screened with ESI (targeted MS/MS mode). The LC system comprised of Phenomenex Kinetex C18 (100 mm × 2.1 mm, 1.7 μm) column. The mobile phase was compose of solvent A (60% ACN+40% H_2_O+10 mM HCOONH4) and solvent B (10% ACN+90% isopropyl alcohol+10 mM HCOONH4) with a gradient elution (0–10.5 min, 30%–100% B; 10.5–12.5 min, 100% B; 12.5–12.51 min, 100%–30% B; 12.51–16 min, 30% B). The flow rate of the mobile phase was 0.3 ml/min. The column temperature was maintained at 50°C, and the sample manager temperature is set at 10°C. Mass spectrometry parameters in ESI+ and ESI− mode are listed as follows: ESI+: Heater Temp 350°C; Sheath Gas Flow Rate, 35 arb; Aux Gas Flow Rate, 15 arb; Sweep Gas Flow Rate, 1 arb; Spray Voltage, 3.0 kV; Capillary Temp, 350°C; S-Lens RF Level, 50%. ESI−: Heater Temp 350°C; Sheath Gas Flow Rate, 35 arb; Aux Gas Flow Rate, 15 arb; Sweep Gas Flow Rate, 1 arb; Spray Voltage, 2.8 kV; Capillary Temp, 350°C; S-Lens RF Level, 50%. At the beginning of the sequence, we run four quality control (QC) samples to avoid small changes in both chromatographic retention time and signal intensity. The QC samples are also injected at regular intervals (every ten samples) throughout the analytical run.

The raw data were acquired and aligned using the Compound Discover (3.0, Thermo) based on the m/z value and the retention time of the ion signals. Ions from both ESI− and ESI+are merge and import into the SIMCA-P program (version 14.1) for multivariate analysis. A Principal Components Analysis (PCA) was first used as an unsupervised method for data visualization and outlier identification. Supervised regression modeling was then performed on the data set by use of Partial Least Squares Discriminant Analysis (PLS-DA) or Orthogonal Partial Least Squares Discriminant Analysis (OPLS-DA) to identify the potential biomarkers. The biomarkers were filtered and confirmed by combining the results of the VIP values (VIP>1.5), t-test (p < 0.05) and fold change (FC>2). The quality of the fitting model can be explained by R2 and Q2 values. R2 displays the variance explained in the model and indicated the quality of the fit. Q2 displays the variance in the data, indicating the model’s predictability.

To compare the lipidomic profiles of our *in vitro* model with those of the IPF patients, we collected the lipidomic data from an IPF study with 18 controls and 22 IPF patients (57). We only looked at the five major lipid classes (P-Glycerol, Glycerolipid, Steroid, Sphingolipid, and Fatty acid) that were both measured in our and the Yan et al. data. In each dataset, the measured intensity of the lipids was first added up according to the lipid classes, then scaled to zero variance and unit variance for the normalization purpose. We then use the two-sample t-test to test for the intensity difference between control and treated/IPF conditions.

### Statistical analysis

Statistics were performed using GraphPad Prism 8.0 software (GraphPad, San Diego, CA). Two-group comparisons were analyzed by unpaired Student’s t-test, and multiple-group comparisons were performed using one-way analysis of variance (ANOVA) followed by Tukey post hoc analysis. Statistical significance was achieved when P < 0.05 at 95% confidence interval.

## Results

### Chronically injured AE2 cells exhibit senescence-like characteristics

Multiple approaches have been utilized for understanding the role of AE2 cells in the pathogenesis of IPF, including exposure to bleomycin, asbestos, or silica. However, they fail to resemble changes seen in the lungs of IPF patients, such as cellular senescence, mitochondrial dysfunction, and impaired proteostasis (47). To address the practical limitation of the current *in vitro* IPF models, we developed a chronic injury model in cultured lung epithelial cells using mouse MLE12 cells, an immortalized cell line used to model alveolar epithelial type 2 cells in culture (Figure 1A). Consistent with findings in the IPF lung epithelium, we found that senescence markers p21 and p53 were detected at higher levels in the chronic-injured model compared to controls (p < 0.05) (Figure 1B). Furthermore, protein and transcript levels of other senescence markers, including γ-H2AX and SASP (senescence-associated secretory phenotype) such as Il-6, Mcp1, TNF-α, and IL-1β were greater in the chronic-injured model (p <0.05) (Figure 1C-D). Moreover, these changes were associated with marked upregulation in senescence-associated beta-galactosidase (SA-βgal) activity in the cells treated with bleomycin and barely evidenced in the control group (Figure 1E). In sum, this model recapitulates some of the features of IPF lung epithelium with prominent AE2 cell senescence.

**Figure 1.**
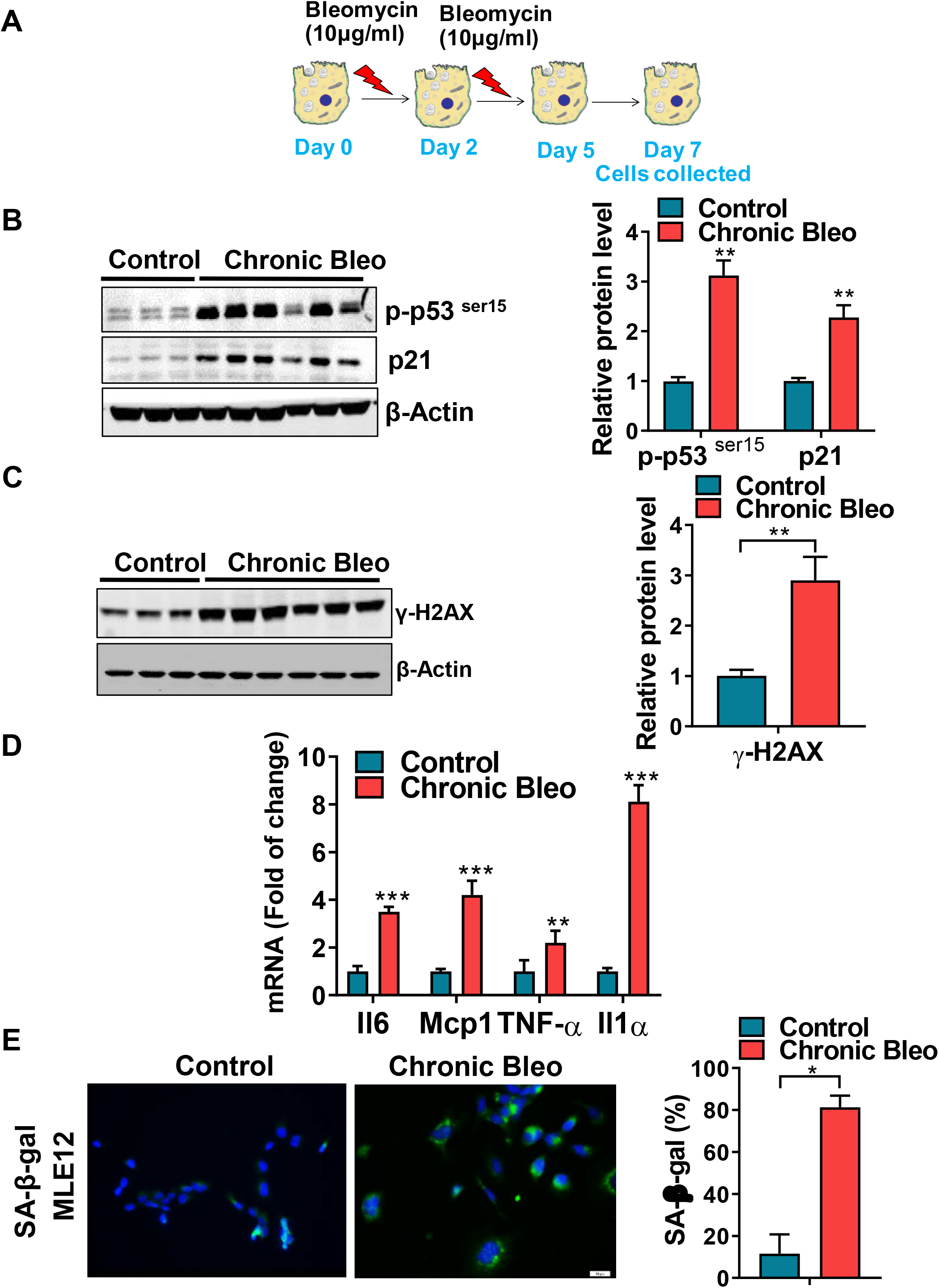
Cellular senescence markers are increased in MLE12 cells exposed to chronic low dose bleomycin. A) Schematic showing the experimental design: On day 0, MLE-12 cells were treated with 10 μ/ml bleomycin for 24h. Day 2, bleomycin was removed, and the culture medium was refreshed. Day 4, cells were treated again with 10 μg/ml bleomycin. Day 5, bleomycin was subsequently removed, and cells were collected at day 7. B) Western blot (WB) for p-p53^ser15^ and p21 in chronically injured lung epithelial cells (with β-actin loading control). Densitometry is shown on the right. C) WB for γ-H2AX in control and bleomycin injured cells (with β-actin loading control). Densitometry is shown on the right. D) Transcript levels for Il6, Mcp1, Tnf-α, and Il1α in controls vs. chronically injured lung epithelial cells. E) SA-β-gal activity (green fluorescence cytoplasmic staining) in control and chronically injured MLE12 cells. Number of SA-β-gal positive cells per 100 cells counted (right). WB images are representative of two different blots. Statistical significance was assessed by Student t-test * p< 0.05, **P < 0.01, ***P < 0.001 versus control group, n=6.

### Mitochondrial function is impaired in MLE12 cells exposed to chronic low dose bleomycin

Previous studies have shown that mitochondrial dysfunction is associated with cellular senescence (8, 31). To verify this in our study, we first assessed whether mitochondrial respiration was decreased in chronically injured AE2 cells using the Seahorse Bioanalyzer to measure the mitochondrial oxygen consumption rate (OCR). As evidenced in Fig 2A, we found that basal respiration, spare respiratory capacity, and ATP-production were significantly lower in the cells treated with low doses of bleomycin (p <0.05). Furthermore, we detected that the extracellular acidification rate (ECAR) was decreased in our chronic injury model as compared to control (Figure 2C), suggesting that the lack of energy production was not being compensated with glycolysis. Consistent with this decline in mitochondrial function and with bleomycin’s known genotoxic effects, we found that chronic exposure to bleomycin readily induced mitochondrial DNA damage and increased mitochondrial ROS levels in MLE12 cells (Fig.2C-D) in MLE12 cells, suggesting a clear link between the presence of cellular senescence markers and mitochondrial dysfunction.

**Figure 2.**
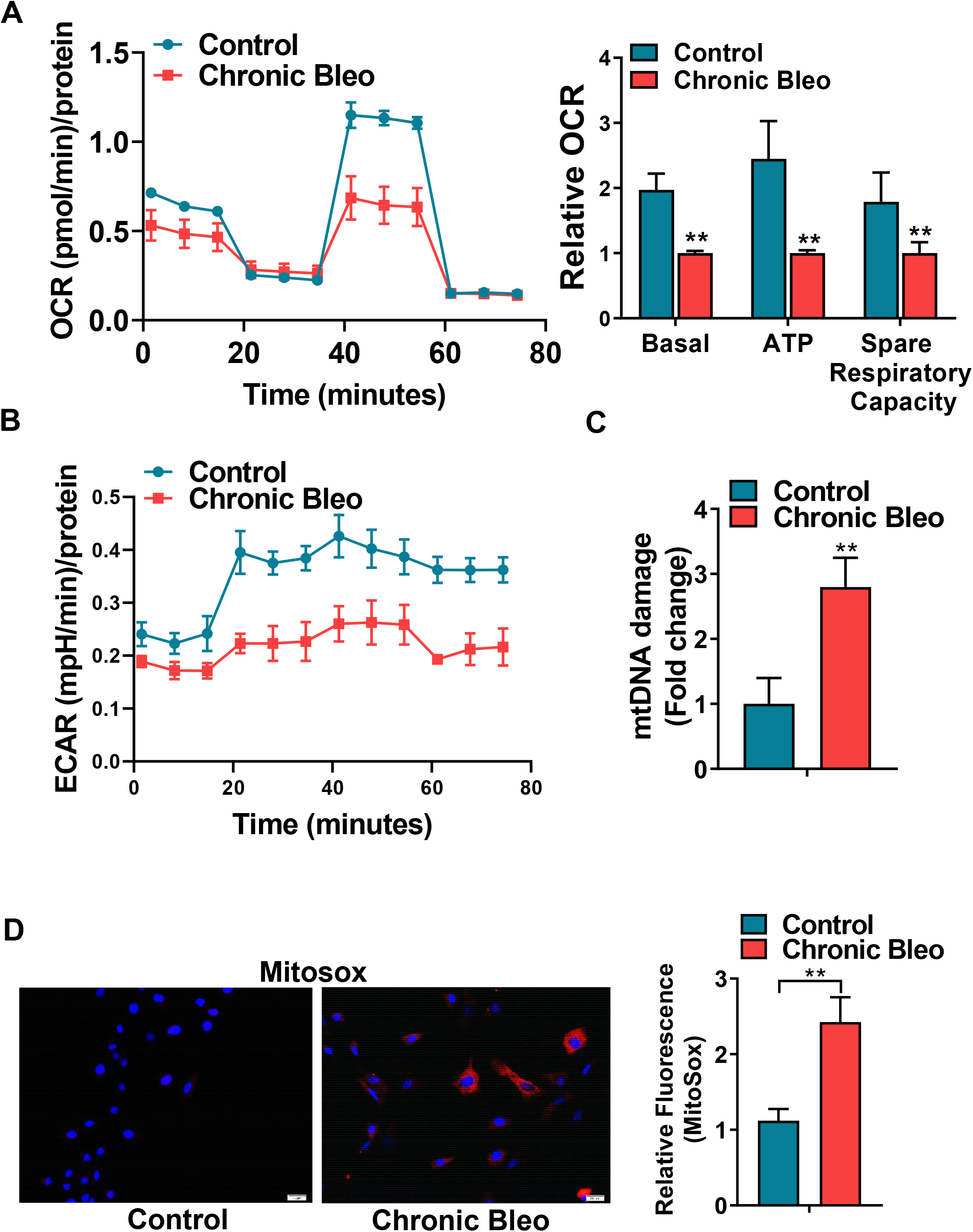
Chronic low dose of bleomycin decreases mitochondrial oxygen consumption in MLE12 cells. A) Basal oxygen consumption rate (OCR), ATP production and spare respiratory capacity in control and chronically injured MLE12 cells. B) Extracellular acidification rate (ECAR) values in control and bleomycin injured cells. C) mtDNA damage quantification in control and bleomycin injured cells. D) Quantification of reactive oxygen species (ROS) by MitoSox staining in control and chronically injured MLE12 cells. Seahorse data are representative of 3 separate experiments. Statistical significance was assessed by Student’s t-test. **P < 0.01 vs. control group, n=6.

### Chronic exposure to bleomycin impaired cellular proteostasis in AE2 cells

Proteostasis failure and accumulation of damaged proteins have been described in pulmonary fibrosis (38). In order to evaluate whether proteostasis network was altered in our chronic injury model, we first used a recently established molecular rotor to measure intracellular protein aggregation. As expected, we found dramatically increased intracellular protein aggregation in chronically injured MLE12 cells compared to controls, which had virtually no aggregated proteins (Figure 3A). Because the ubiquitin-proteasome system (UPS) represents the major regulator of protein homeostasis in eukaryotic cells by degrading misfolded proteins, we assessed whether the UPS activity and gene levels of key proteasome subunits were perturbed in chronically injured MLE12 cell. Consistent with an increase in protein aggregation, we found that chymotrypsin-like, trypsin-like, and caspase-like activities were significantly reduced when comparing levels in injured cells versus controls (Figure 3B-D). Next, in order to evaluate whether the reduction in 26S proteasome activity was due to impaired proteasome subunit production, we evaluated the expression of proteasome subunit. As shown in Figure 3E, we found that expression of the subunits Psmd12, Psmd5, PsmbB6/β1 were markedly decreased when comparing chronically injured to control MLE12 cells, indicating that the decline in the proteasome function is mediated by downregulation in proteasome subunits expression. Taken together, our data suggested that the decline in molecular proteostasis seen in chronically injured MLE12 cells might be linked to dysregulations in the activity of the proteasome.

**Figure 3.**
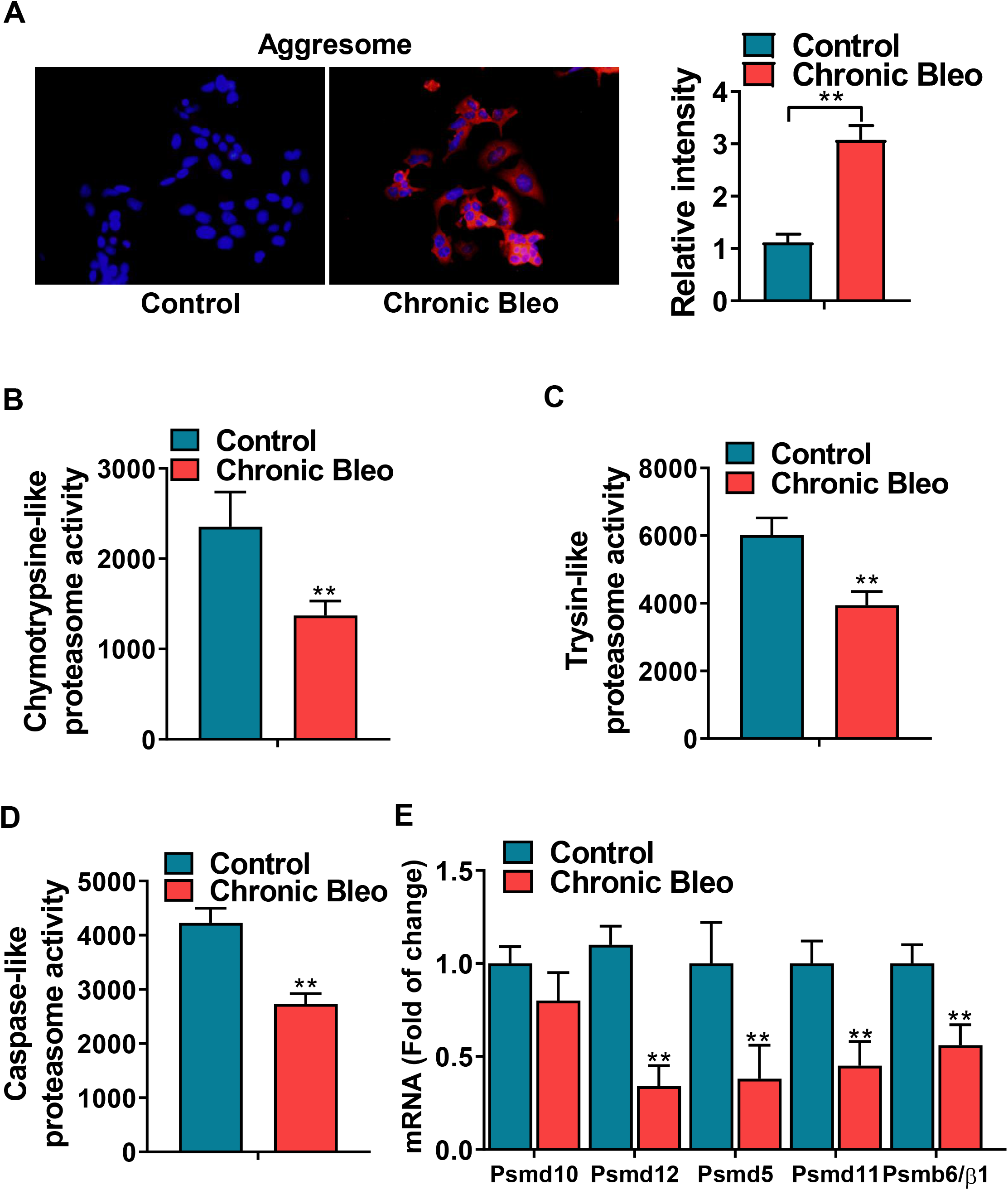
Cellular proteostasis declines in chronically injured lung epithelial cells. A) Proteostat staining (aggresome) in controls and chronically injured MLE12 cells. Quantification is shown on the right. B, C, D) Chymotrypsin-like, trypsin-like and caspase-like activities in control and bleomycin injured MLE12 cells. E) Transcript levels for PSMD10, PSMD12, PSMD5, PSMD11, and PSMB6 in controls vs. chronically injured MLE12 cells. Statistical significance was assessed by Student t-test ** p< 0.01 versus control group, n=6.

### HSF-1 transcriptional activity is impaired in chronically injured MLE12 cells

We next evaluated whether heat shock proteins (HSPs) network was altered in chronically injured MLE12 cells. We focused on HSPs expression since the regulation of these chaperones is believed to be the major contributor of cytoplasmic proteostasis. Consistent with the misfolded protein accumulation, we found that gene expression of *Hsp70, Hsp90, and Hsp40* was significantly decreased in chronically injured MLE12 cells (Figure 4A). Unexpectedly, as shown in Fig 4B, we only found a difference in the protein levels of HSP70. We next sought to determine whether the downregulation of this pathway was associated with a decrease in the levels of the heat shock factor 1 (HSF1), a transcription factor that regulates the expression of HSPs to maintain cellular proteostasis. As expected, we found that nuclear levels of HSF1 were decreased in chronically injured MLE12 cells. Since, HSF1 functional activity can be regulated by post-transcriptional modification, including phosphorylation at serine 307 and phosphorylation at serine 326, we next thought to investigate whether this reduction in the expression of the cellular chaperone was due to post-transcriptional inactivation of the HSF1. Remarkably, p-HSF1^*Ser307*^ expression was increased by over 2-fold in chronically injured MLE12 cells, while expression of p-HSF1^*Ser326*^ was found downregulated in our chronic model. (Fig. 4C). Thus, our data suggest that HSF1 transcriptional activity is impaired possibly due to inhibited phosphorylation of the Ser326, a required step in the induction of the factor’s transcriptional competence, suggesting a link between impaired HSF1 transactivation, reduction in proteins chaperones, and proteostasis collapse. Altogether, these findings indicate that cytoplasmic proteostasis is profoundly altered in chronically injured MLE12 cells, perhaps due to combined effects on the expression and clearance of the chaperone of misfolded proteins from cells.

**Figure 4.**
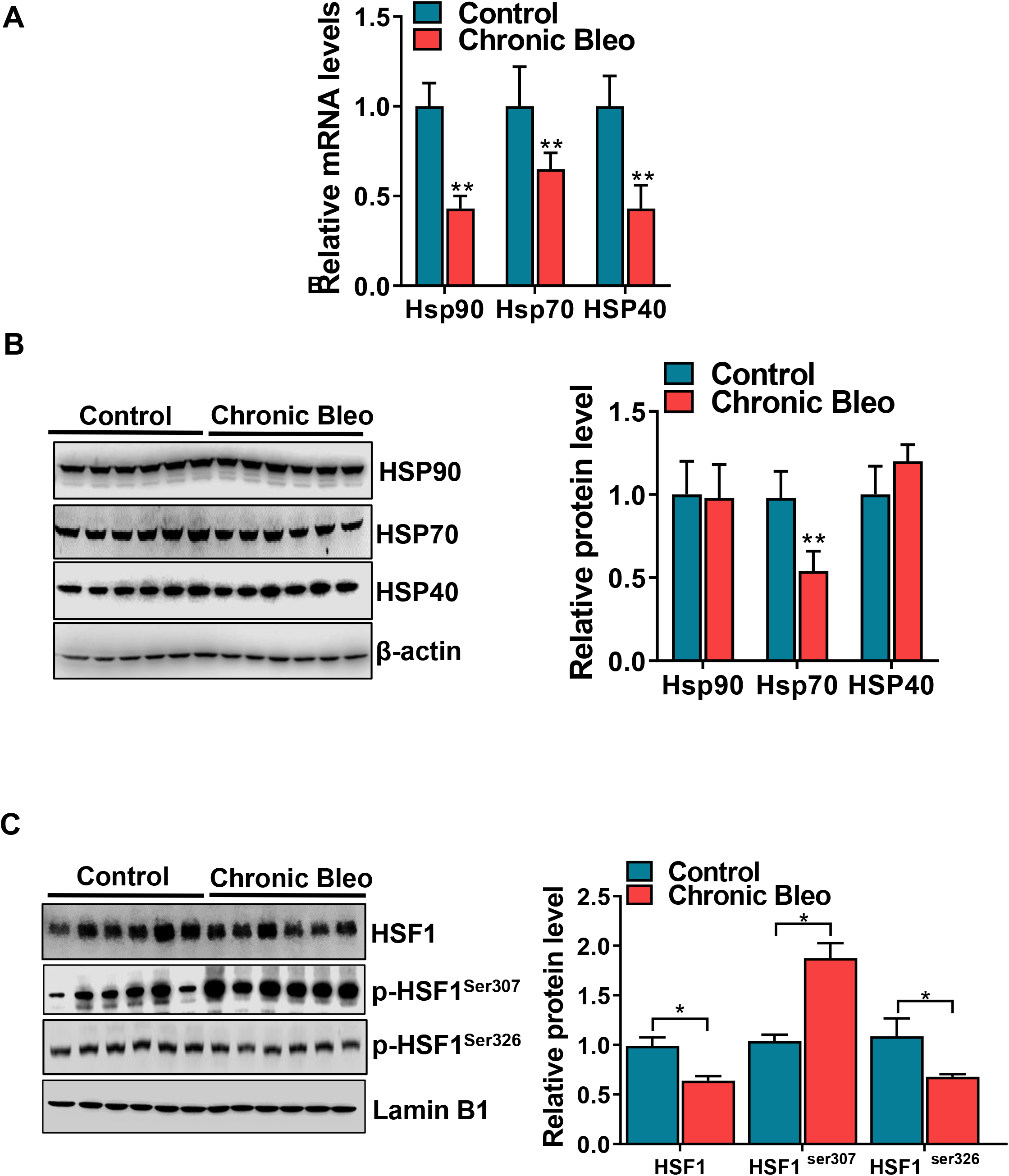
The HSF1 activity is impaired in MLE12 cells exposed to chronic low dose bleomycin. A) Transcript levels for Hsp90, Hsp70, and Hsp40 in control and bleomycin injured MLE12 cells. B) WB for HSP90, HSP70 and HSP40 in chronically injured MLE12 cells (with β-actin loading control). Densitometry is shown on the right. C) WB for HSF1, p-HSF1^ser307^, p-HSF1^ser326^ in control, and bleomycin injured MLE12 cells (with Laminin-B1 loading control). Densitometry is shown on the right. Western blot images are representative of two different blots, and results of densitometry analysis are depicted in bar graphs (n= 6, per group). Statistical significance was assessed by Student t-test * p<0.05, ** p< 0.01 versus control group, n=6.

### RNA-seq analysis of differentially expressed genes in bleomycin injured AE2 cells and controls

To better understand these cellular stress pathways in chronically injured MLE12 cells, we performed RNA sequencing (RNA-seq) to analyze the transcriptomes of these cells. Approximately 25 million reads (94% of clean reads) mapped to the reference genome. The expression of each transcript in each sample was measured as the expected number of fragments per kilobase of transcript sequence per millions of base pairs sequenced (FPKM). Genes with values of FPKM > 1 were considered to be genes expressed in this study. The dynamic range of the expression values was calculated and is presented as a Boxplots of log10-transformed for each replicate (Figure S1A), and the FPKM density distribution is shown in Figure S1B. Overall, the transcriptome data were highly reproducible with a correlation coefficient of >90% across all samples and >95% between biological replicates. These results indicated that our RNA-seq data was reliable and reproducible. The principal component analysis (PCA) showed a clear separation of gene expression between the two populations, controls versus chronically injured MLE12 cells (Figure 5A). As shown in the volcano plot, we identified 8484 differential expressed genes, including 4777, with increased and 3707 with reduced expression, representing a robust transcriptional response (Figure 5B). In addition, 558 and 139 unique differentially expressed genes (DEGs) were identified in bleomycin injured and control MLE12 cells, whereas 10777 DEGs were overlapped between the two libraries (Figure 5C). Heatmap representing a 2D hierarchical clustering of genes shows differential expressions in the two groups (Figure 5D).

**Figure 5.**
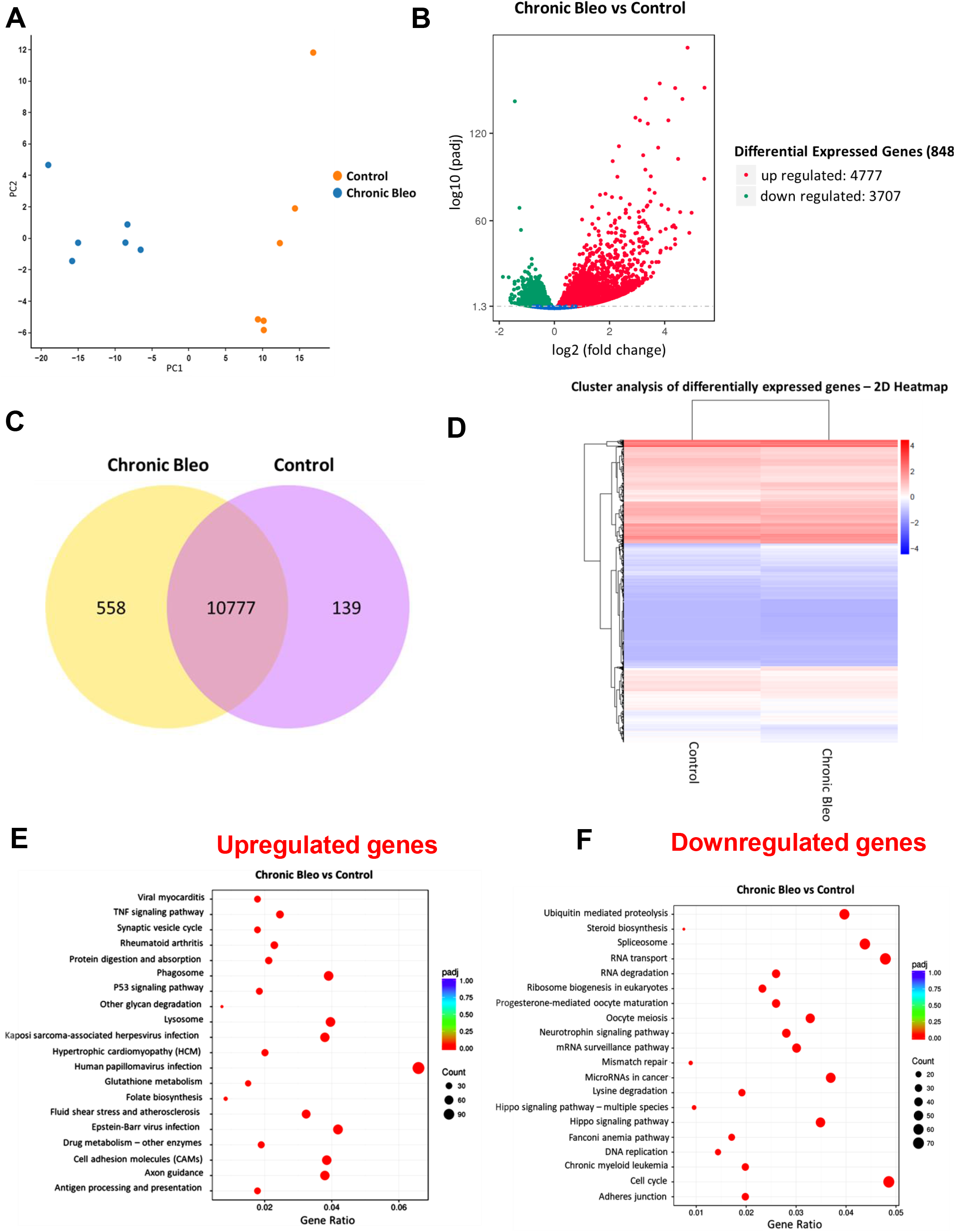
Transcriptome signature of controls and chronically injured MLE12 cells. A) Principal component analysis (PCA) of RNA-seq data of control and bleomycin injured MLE12 cells (n = 6). B) Volcano plot of differentially expressed genes of the bulk RNA-seq data from controls and chronically injured MLE12 cells. C) Venn diagram depicting the comparison and overlap of differentially expressed genes in controls and chronically injured MLE12 cells. D) Heatmap representing 2D hierarchical clustering of genes, showing differentially expressed genes in control and bleomycin injured cells. KEGG pathway enrichment analysis of predicted biological processes and genes induced in bleomycin injured cells. Advance bubble chart shows enrichment of differentially expressed genes in signaling pathways (E) upregulated and (F) downregulated. Y-axis label represents the pathway, and the X-axis label represents gene ratio. The size and color of the bubble represent the amount of differentially expressed genes enriched in pathway and enrichment significance, respectively.

Gene Ontology (GO) enrichment analysis of differentially expressed genes was implemented by the cluster profiler R package. In total, 4159 up-regulated DEGs were annotated in 686 significant GO terms, most of which were involved in biological processes related to the lysosome (184), wound healing (136), extracellular matrix, (118), aging (113), and response to IFN (33) (Figure 5E). Also, a total of 3316 down-regulated DEGs were significantly annotations involved mRNA processing (221), histone modification (203), DNA repair (196), and RNA splicing (179) (Figure 5F). We next performed a Kyoto Encyclopedia of Genes and Genome (KEGG) pathway enrichment analysis to annotate the functions of the DEGs identified in the chronically injured MLE12 (Figure 5E). We identified a total of 7002 genes that were differentially expressed, of which 1792 were up-regulated, and 1462 were down-regulated. Notably, the up-regulation of key pathways were identified, including PI3K/AKT (108), MAPK-signaling (96), cell adhesion molecule (69), TNF signaling pathway (44), and p53 signaling pathway (33(Figure S2A). Downregulated DEGs were mainly significantly enriched in the cell cycle (71), ubiquitin-mediated proteolysis (58), lysine degradation (28) and steroid biosynthesis (11) (Figure S2B). For Reactome analysis, the top 20 most significantly up-regulated and down-regulated genes are shown in Figure S3A-B. The complete list of GO terms, KEGG pathways, and Reactome are listed in Supplementary File X.

Next, in order to evaluate whether the transcriptome data from our chronic model shares similarities in gene expression with previous reports on IPF lungs, we conducted a principal component analysis (PCA) to compare the overall transcriptomic profiles in our data to those from Xu et al. (56) and Nance et al. (32). Even though the samples were generated in three different studies, the control samples and the bleomycin-treated/IPF samples can be separated using the first two principal components (PCs), implying that the overall transcriptomic profiles of our *in vitro* model resemble those of the actual IPF patients (Figure 6A). By performing the DE analysis, we were able to identify a set of up-regulated and down-regulated genes that demonstrate similar transcriptomic changes between control and IPF conditions and between control and our *in vitro* model. This result suggests that by using chronic bleomycin exposed MLE12 cells, we were able to mimic the transcriptomic alterations of those seen in the lung epithelium of IPF patients. According to our GO enrichment analysis, the up-regulated genes in our treated cells or IPF patients are responsible for the development, differentiation, and proliferation processes, including cornification, epidermis development, keratinization, keratinocyte differentiation and skin development (Figure 6B and Table supplementary 1A). In contrast, the down-regulated genes are involved in multiple biosynthetic and metabolic processes, including cholesterol biosynthetic process, sterol metabolism, regulation of lipid metabolism process, and secondary alcohol metabolism (Figure 6C and Table supplementary 1B). Thus, these results established that our chronic-injury model shares similarities in gene expression with previous reports on IPF lungs.

**Figure 6.**
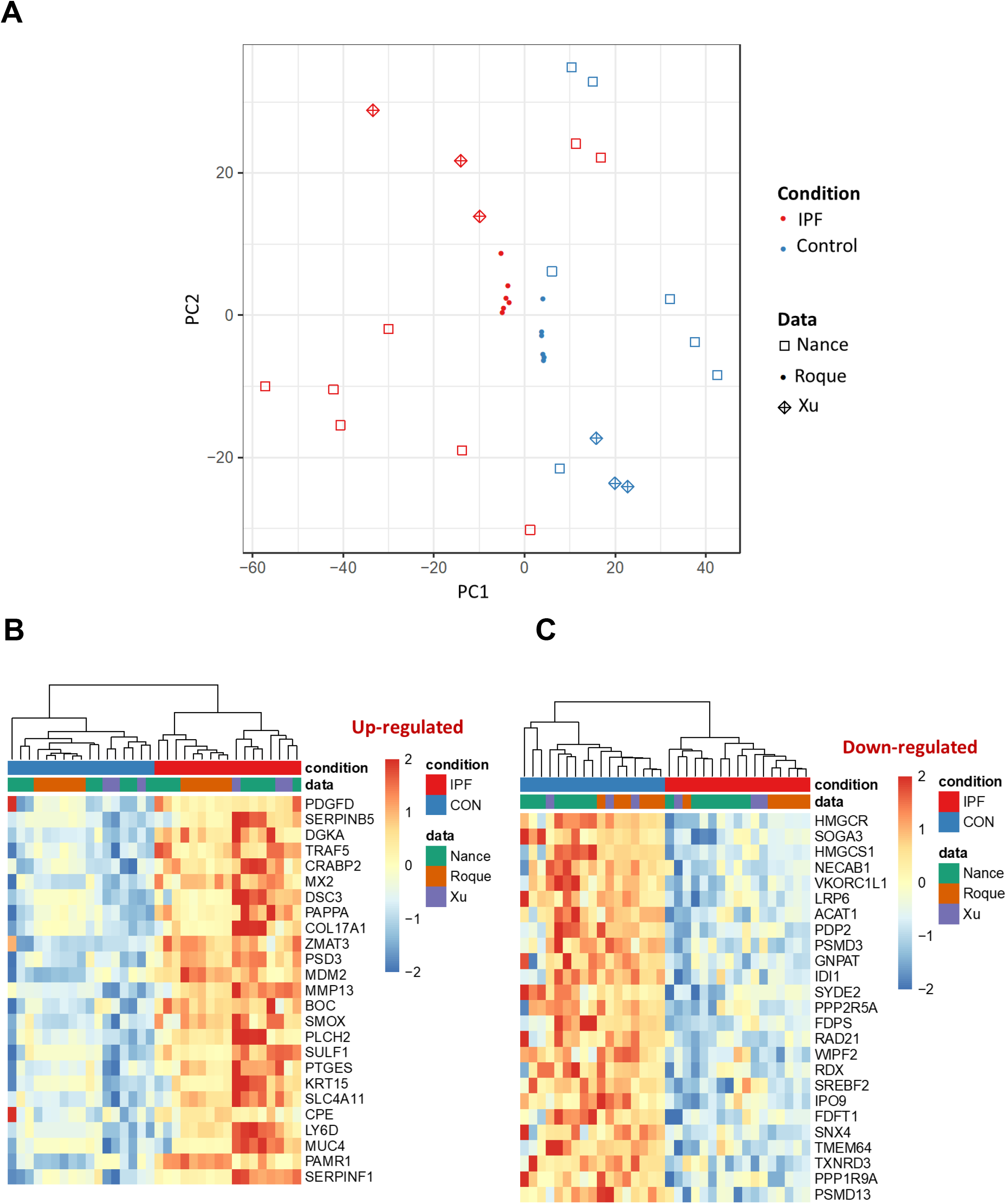
Comparison of transcriptomic network changes in chronically injured MLE12 cells versus IPF lung epithelial cells. A) Principal component analysis (PCA) of transcriptomic profiles comparison from different datasets of IPF, bleomycin injured MLE12 cells, and controls. B) Dendogram cluster of significant shared changes in gene expression in chronically injured MLE12 cells and IPF lung epithelial cells. Red indicates relative higher expression, blue mean relative lower expression.

### Metabolomics characterization of bleomycin injured MLE12 cells

We next examined whether other characteristics of IPF lung were evident in our chronic model. To this end, deficient AE2 cell metabolism has recently emerged as an essential player in the pathogenesis of IPF. To test whether cellular metabolism was altered in chronically injured MLE12 cells, we determined metabolome signature in our cells by LC-MS/MS analysis. We first use principal-component analysis (PCA) to analyze all observations acquired in positive and negative ion modes. PCA score plots showed the noticeable separation trend between the control and bleomycin groups, indicating that cellular metabolic states of chronically injured MLE12 cells were significantly changed in relative to controls (Figure 7A-B). We then built OPLS-DA and PLS-DA models in order to eliminate any non-specific effects and confirm the metabolites. Both models showed a clear separation of control versus chronically injured MLE12 cells (Figure S4A-D). Next, a 1000-time permutation test was conducted to further validate the OPLS-DA model (Figure S5A-B). The empirical p values for R2Y (p<0.01) and Q2 (p<0.001) indicate that the observed statistic is not part of the distribution formed by those from permuted data. Thus, the clustering of the data is statistically significant. An S-plot was then constructed to visualize the loading of the POLS-DA model (Figure S5C-D) and determine which variables are the best discriminators between the different groups. Afterward, the significantly changed metabolites between the groups were filtered out based on VIP values (VIP>1.5), and Figures S6A-B show the distribution of numerical values. The PLS-DA loading plot is showed in Figure S6C-D, and the metabolites with the red box are labeled as significant compounds. These results illustrated that the metabolites in the control and bleomycin treated groups had been completely separated, either the positive or the negative ion mode.

**Figure 7.**
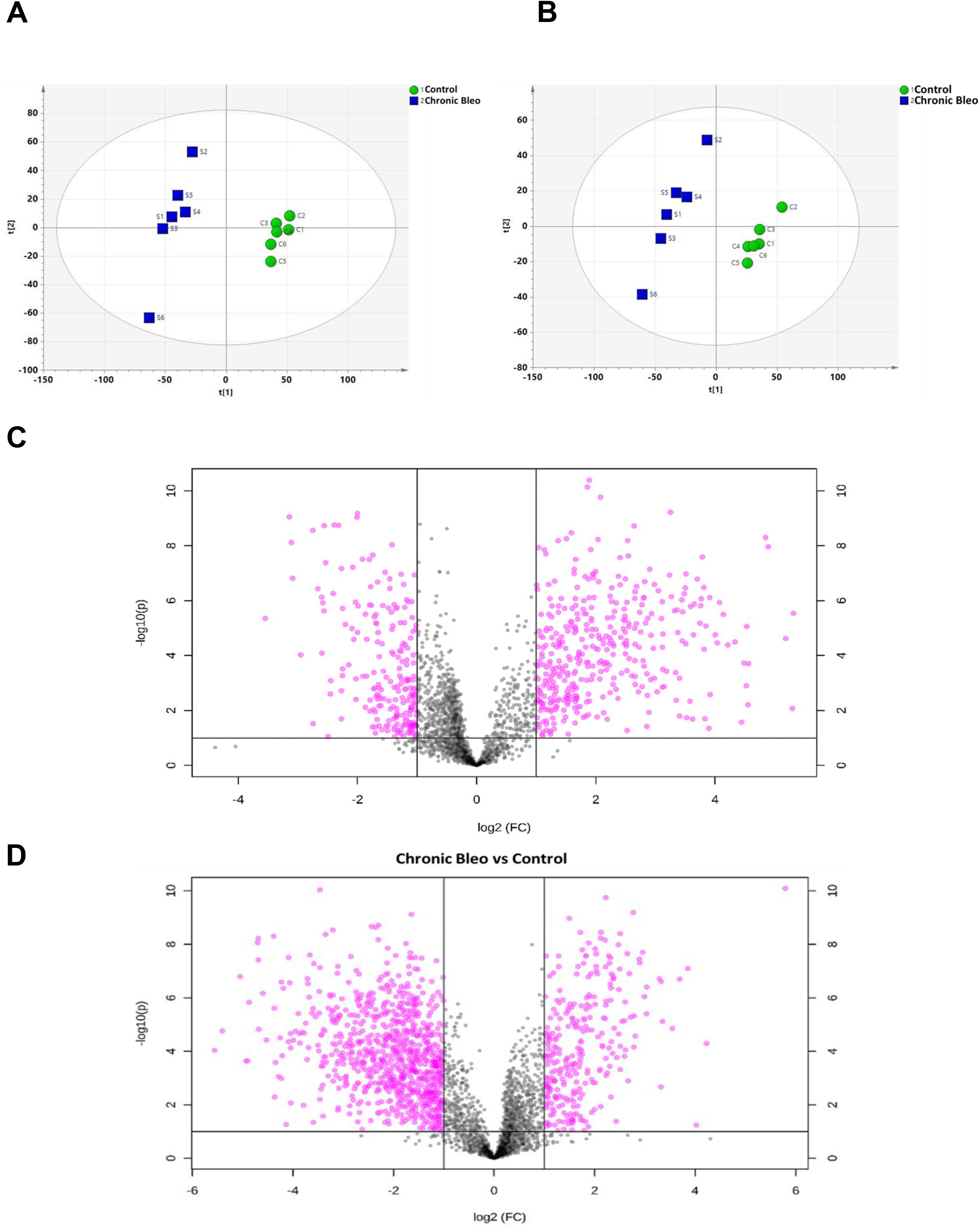
Non-targeted metabolomics analysis between controls and chronically injured MLE12 cells. A) Principal component analysis (PCA) of the metabolomic profile of controls and injured cells (positive ion mode). B) PCA score plot of negative ion mode. Volcano plot of differentially expressed metabolites from control and bleomycin injured MLE12 cells with (C) positive and (D) negative mode.

To screen the differential metabolites, the VIP in the OPLS-DA model (VIP>1.5), FC > 2.0, and p-value of Student’s t-test (p<0.05) were used as the criteria. The results of screening differential metabolites were visualized in the form of volcano plots (Figure 7C-D). Each point in the volcanic map represents a metabolite. A total of 754 metabolites (positive ion mode) and 441 metabolites (negative ion mode) were significantly changed after bleomycin exposure. Relative to the control group, 182 metabolites were significantly upregulated, and 122 were significantly downregulated, including 153 amino acids and their derivatives, 11 glycerophospholipids, 31 nucleotides, and their derivatives, and others. Hierarchical clustering analysis was then performed to study the metabolomic profiles between control and bleomycin injured groups. The heat map representation of metabolome profiles analyzed by hierarchical clustering analysis is summarized in Figure 8A-B (with complete data in Supplementary information). Differential metabolites were identified using the KEEG Metabolome Database, and the results are shown via a bubble plot (Figure 8C-D). Pathway impact values indicate that the main enrichment metabolic pathways of differential metabolites after bleomycin injury included reduced sphingolipid metabolism, fatty acid metabolism, gluconeogenesis, glutamate, and tyrosine metabolism. In addition, we found upregulation of phospholipid biosynthesis, urea cycle, and malate-aspartate shuttle. Next, we compared the metabolomics profiles of our *in vitro* model with those of the IPF patients (18). Our metabolomic analysis identified six metabolite classes (glucose, phenylalanine, proline, succinct acid, threonine, and uracil) that undergo similar abundance changes in the alveolar epithelial cells with bleomycin administration compared with those observed in the lung epithelium of IPF patients (Figure 9). Among these metabolite classes, only glucose demonstrate reduced intensity in treated cells compared with the control cells, and the other five classes all have increased intensity in the treated cells.

**Figure 8.**
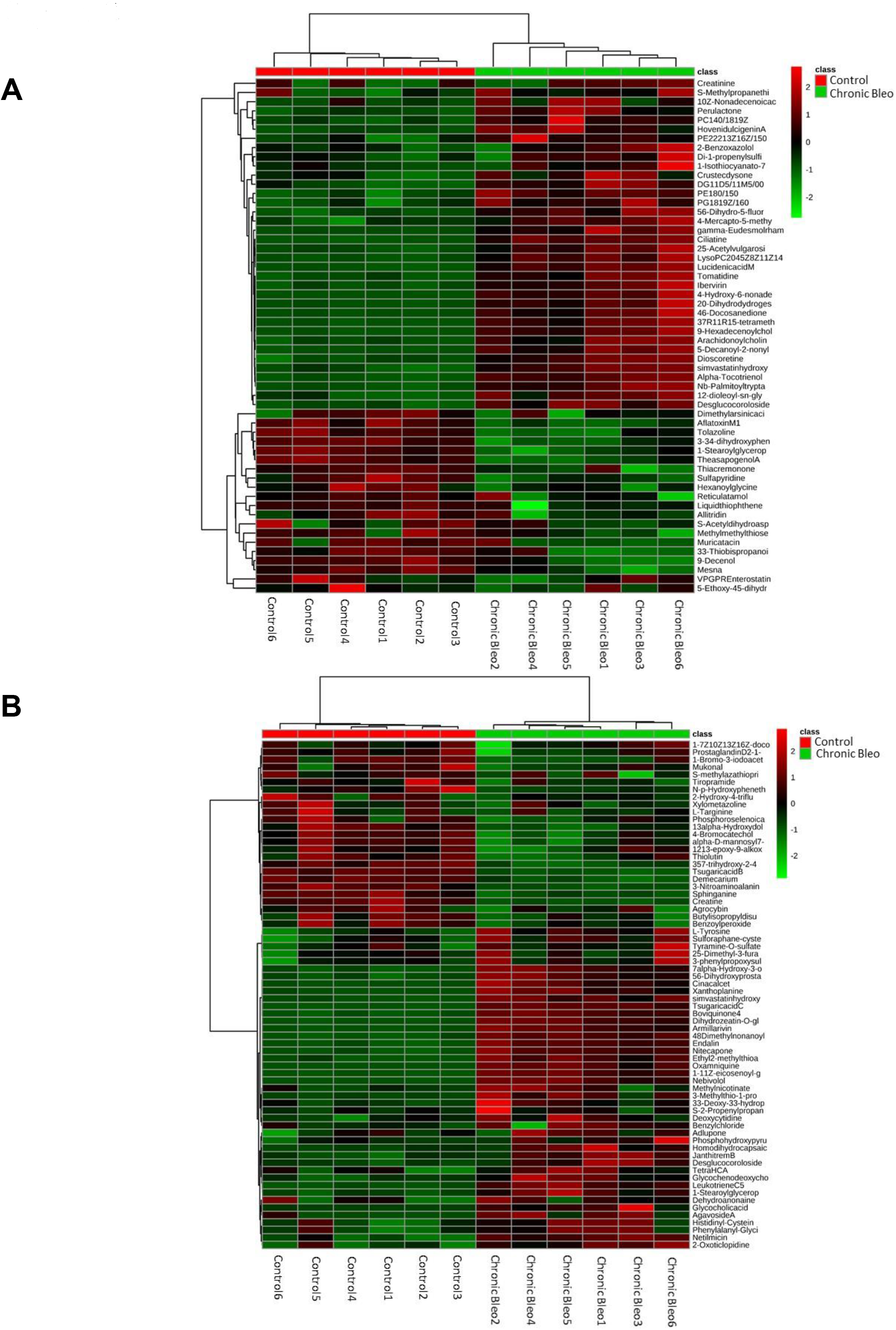
Heatmap showed the differences of metabolites between the control group and bleomycin injured group. A) Heatmap representing 2D hierarchical clustering of metabolites, showing differentially expressed metabolites in control and chronically injured MLE12 cells. Red indicates relative higher expression, green mean relative lower expression.

**Figure 9.**
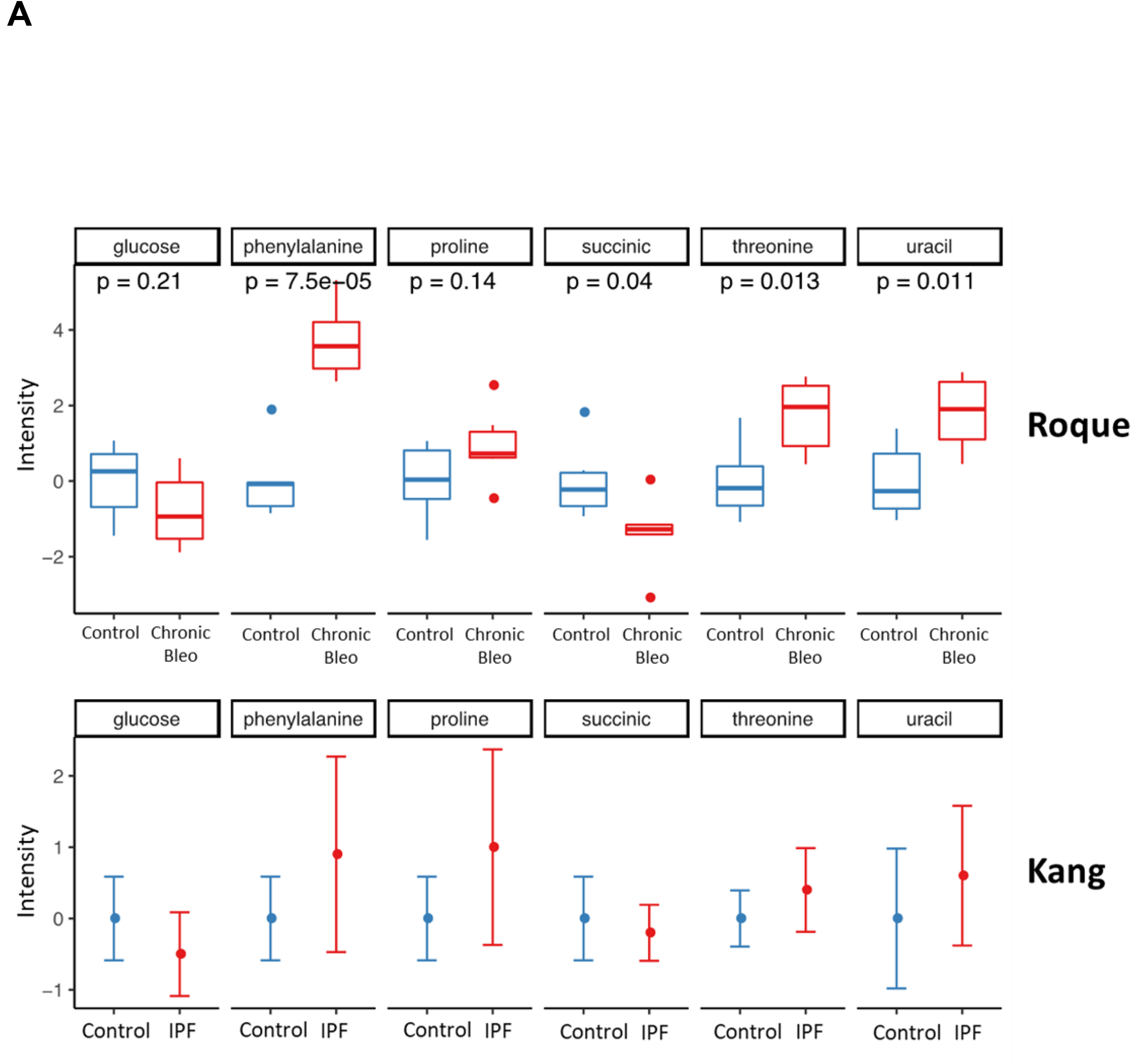
Shared chronically injured MLE12 cells and IPF lungs metabolites network. Heatmap showed the differences of metabolites between the control group and bleomycin injured group. A) Metabolite intensity comparison with a different dataset of IPF vs. control and bleomycin injured MLE12 cells vs. control.

### Multivariate analysis of Lipidomic profiling of chronic-injured AE2 cells

Alterations in lipid metabolism are one of the most prominent metabolic changes observed in the lung epithelium of IPF patients. In order to investigate the global lipid variations in our *in vitro* model, we began grouping the samples using a PCA score plot, comparing controls and chronically injured MLE12 cells. As shown in PCA plots (Figure 10A-B), an overview of all samples in the data can be observed and exhibit a clear grouping trend between the two groups. PLS-DA and OPLS analysis (Figure S7A-D) was performed using the data from UPLC-MS; a clear separation of control versus bleomycin injured groups was observed. Afterward, the significantly changed metabolites (positive and negative ions) between the groups were filtered out based on VIP values (VIP > 1.5), and Figure S8A-B shows the distribution of numerical values. The PLS-DA loading plot is showed in Figure S8C-D, and the metabolites with the red box are labeled as significant metabolites. After obtaining the raw intensity data, average group intensities, and fold-changes, we selected the metabolites with VIP>1.5, FC>2.0, and p<0.05 as significant compounds. Univariate analysis, including fold change and t-test, was performed on Volcano plot (Figure 10C-D). A total of 1302 lipid species (positive ion mode) and 901 lipid species (negative ion mode) were significantly changed after bleomycin exposure. The chemical structures of important metabolites were then identified according to the Human Metabolome Database, Metlin and The Mass Bank using the data of accurate masses and MS/MS fragment and showed in Supplementary information X. To summarize the univariate results, we performed a hierarchical cluster analysis (HCA) of lipidomic data from control and chronically injured MLE12 cells by using the complete linkage algorithm of the program Cluster 3.0 (Stanford University) and the results are visualized using Treeview (Stanford University). Metabolite ratios from two independent experiments of every significant lipid species were used for HCA. Color intensity correlates with the degree of increase (orange) and decreases (blue) relative to the mean lipid ratio in the positive or the negative ion mode (Figure 11A-B). Potential target pathway analysis demonstrated that the lipids identified by statistical analysis are accountable for phosphatidylcholine (PC), triacylglyceride (TG), phosphatidylethanolamine (PE), lysophosphatidylcholine (LPC), phosphatidylinositol (PI), sphingomyelin (SM), phosphatidylserine (PS), phosphatidylglycerol (PG), ceramide (Cer), phosphatidic acid (PA), and dimethylphosphatidylethanolamine (dMePE) (Figure 12A-B). Finally, our lipidomic analysis shows that the major lipid classes undergo similar abundance changes in the alveolar epithelial cells after bleomycin exposed compared with those observed in the lung epithelium of IPF patients (57). Especially, fatty acid, P-glycerol, glycerolipid, and sphingolipid all demonstrated increased intensity in the treated/IPF cells compared with the control cells (Figure 12C).

**Figure 10.**
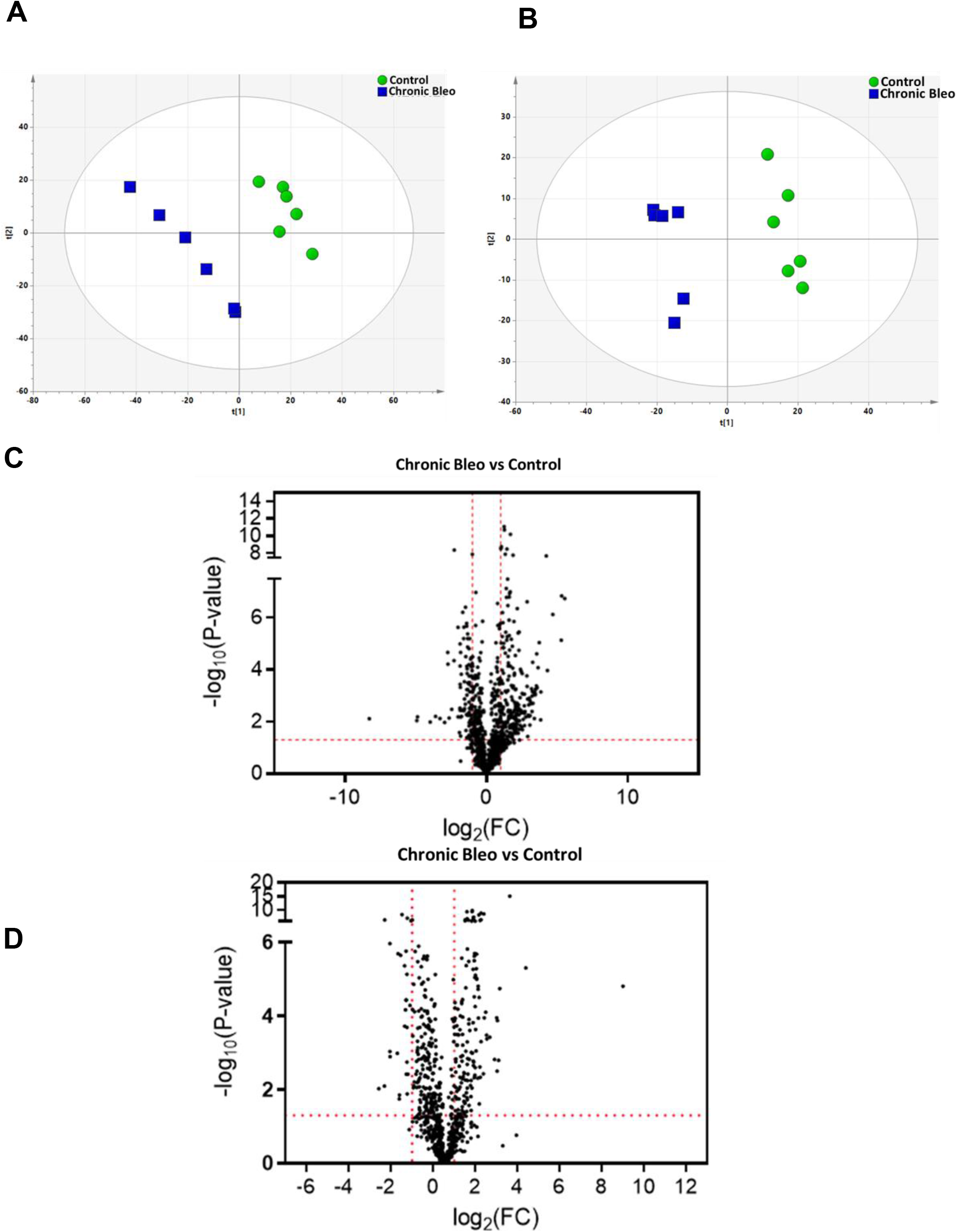
Non-targeted lipidomics analysis between controls and chronically injured MLE12 cells. A) Principal component analysis (PCA) of lipid species profile of controls and injured cells (positive ion mode). B) PCA score plot of negative ion mode. Volcano plot of differentially expressed lipids from control and bleomycin injured MLE12 cells with (C) positive and (D) negative mode.

**Figure 11.**
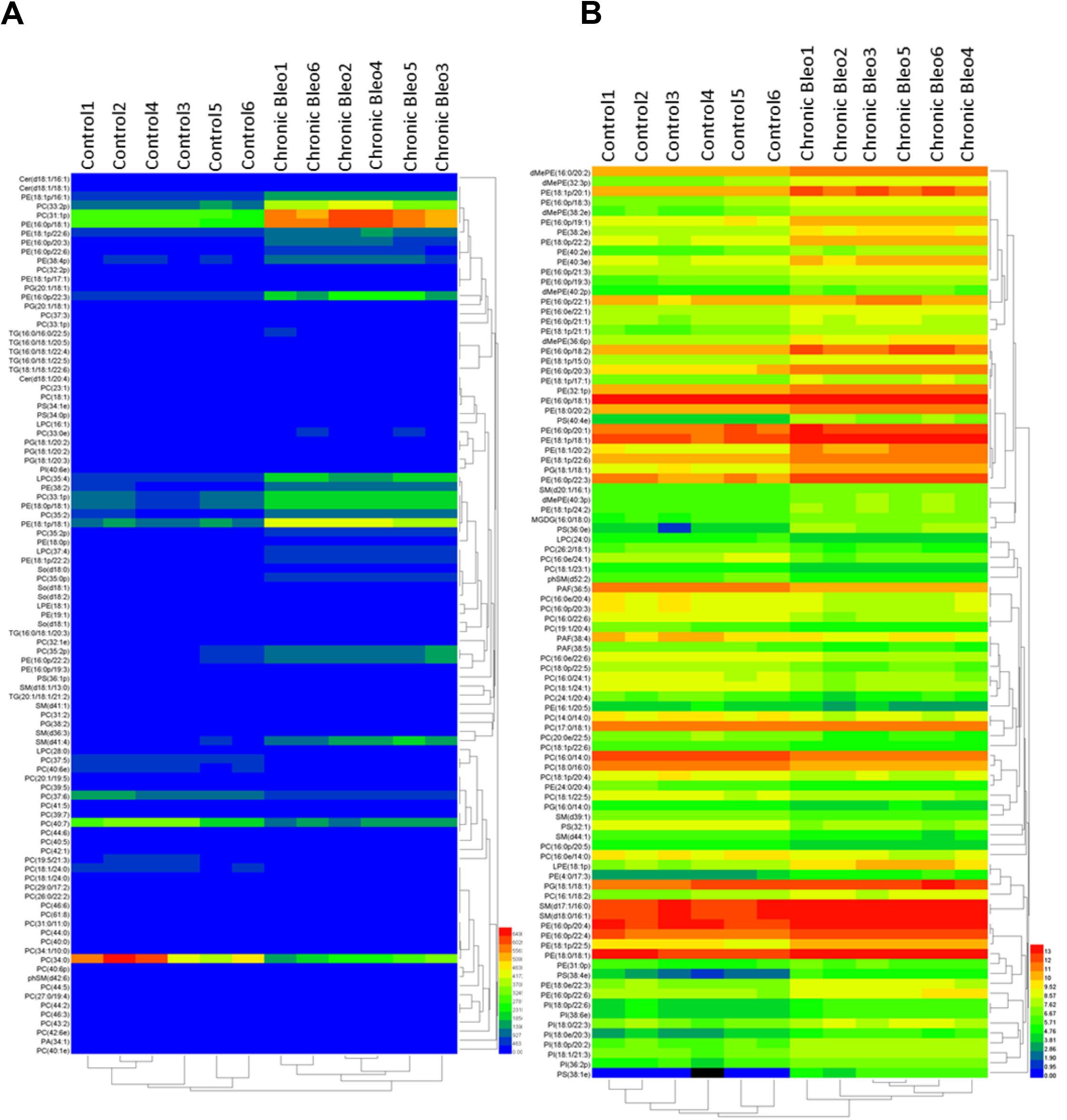
Hierarchical cluster analysis showed the differences of lipids species between the control group and bleomycin injured group. A) Heatmap representing 2D hierarchical clustering of lipids, showing differentially expressed metabolites in control and chronically injured MLE12 cells. Red indicates relative higher expression, blue mean relative lower expression.

**Figure 12.**
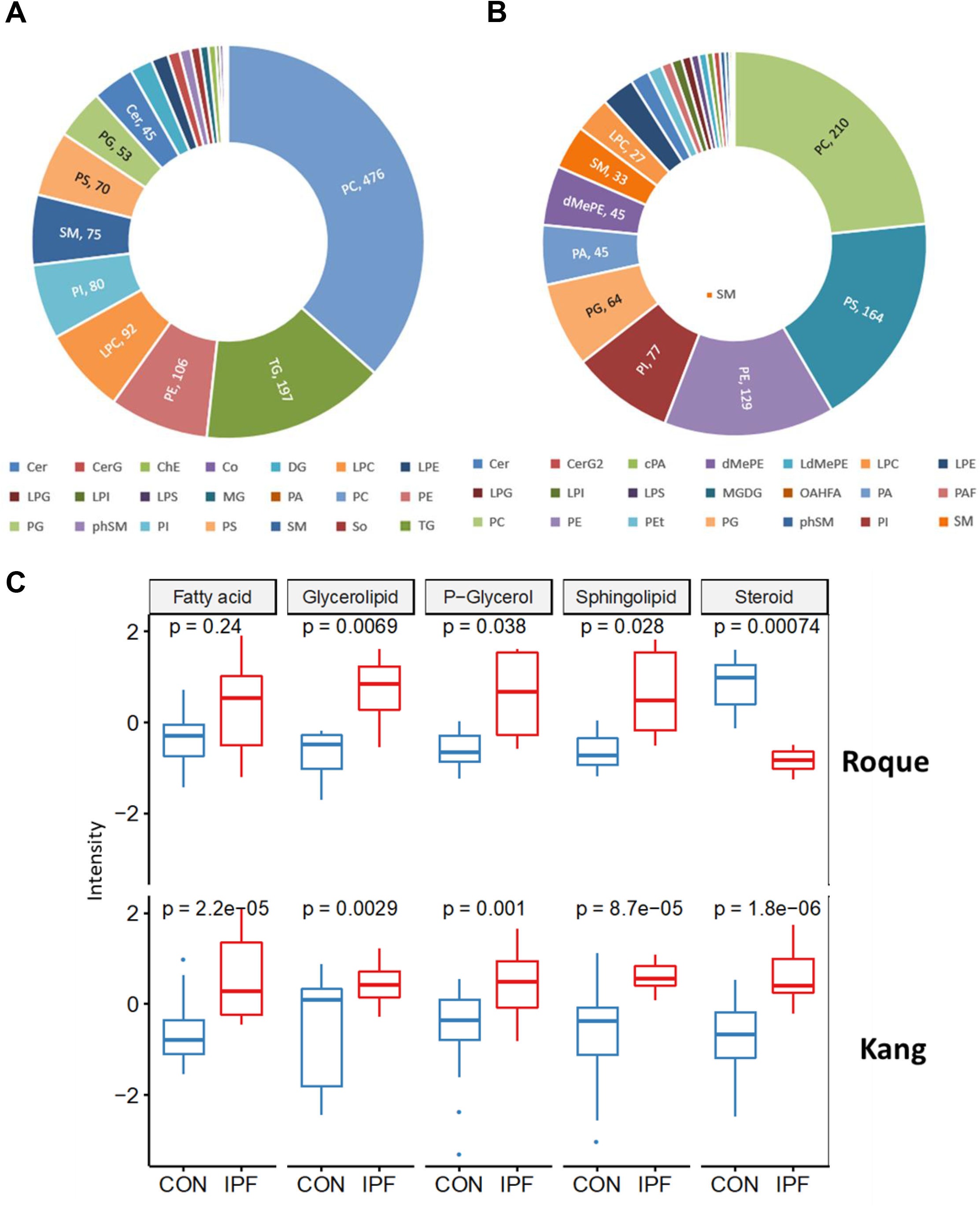
Comparison of lipid pathway enrichment in chronically injured MLE12 cells and IPF lungs. The distribution of the different lipid species in controls and bleomycin injured cells, (A) positive and (B) negative ion mode. C) Metabolite intensity comparison between dataset of IPF lungs and bleomycin injured MLE12 cells vs. control.

## Discussion

The pathophysiology of IPF, an age-related and rapidly progressive lung disease remain poorly understood. Sustained injury to alveolar epithelial type II (AE2) cells leading to aberrant pathways activation has been proposed as a major hallmark of the IPF lung epithelium (6, 33, 44). Up to date, no appropriate *in vitro* models have been developed to explore the cellular mechanism underlying this disease. Here, our objective was to create such model that mimics the molecular and functional characteristics of the IPF lung epithelium.

We demonstrated that chronically injured AE2 cells exhibited all the features of senescence cells, including increases in p21, SASP, and mitochondrial dysfunction. Consistent with prior studies, our model also revealed proteostasis collapse leading to the accumulation of damaged proteins within the cellular cytoplasm secondary to decreases in proteasome activity (26, 35, 48, 50). We found that given this accumulation of protein aggregates, the expression of different heat-shock-proteins was reduced as a result of compromised nuclear translocation of HSF1. Next, to fully characterize our model, we performed a transcriptomic, metabolomic, and lipidomic analysis, which revealed significant similarities to the findings reported in the IPF lung epithelium. In sum, hallmarks of idiopathic pulmonary fibrosis and aging were evidence in our chronic injury model. We believe this model will be ideally suited for use in uncovering novel insights into the gene expression and cellular pathways activation of the IPF lung epithelium and performing high-throughput screening of pharmaceutical compounds.

Combined, the findings presented in our model recapitulate the major elements described in the pathogenesis of IPF. For instance, the presence of senescence phenotype in lung epithelium has emerged as a major contributor to aging and multiple respiratory diseases, including lung fibrosis (3, 24, 28). We reported the presence of cellular senescence markers, such as p21, p53, γ-H2AX and SA-βgal staining, which along with the expression of SASP components represent a source of chronic profibrotic signaling through the activation of Il-6, Mcp1, TNF-α, and IL-1α. The presence of this secretory phenotype has already been confirmed in AE2 cells and lung fibroblasts of IPF patients (2, 54). We previously demonstrated that chronic activation of the mTOR/PGC-1β pathway induces lung epithelial cells into a senescence phenotype, possibly due to augmented ROS-induced molecular damage (51). The expression of SASP has been recognized as another driver of the lung fibroblast to myofibroblast differentiation and a direct contributor to the excessive collagen deposition (21, 30). Our model emulates this major “hallmark” of age-related lung fibrosis, serving as a future tool to fully elucidate the involved regulatory mechanism with the goal of developing therapeutic approaches.

Growing evidence indicates that aging and cellular senescence are tightly interrelated to mitochondrial dysfunction (16, 54). Accumulation of dysmorphic/dysfunctional mitochondria has been described in several aged tissues, including AE2 cells of IPF lungs. In addition, senescent lung fibroblasts from IPF patients have shown diminished oxidative phosphorylation associated with an increased ROS production, possibly perpetuating the senescence phenotype and leading to accumulation of dysfunctional mitochondria (1, 42). In our study, we successfully demonstrated that chronic use of low dose bleomycin in AE2 leads to the findings described above, emulating the prior reported auto-amplifying loop of mitochondrial-derived ROS and senescence. Although our results add up to the free radical theory as a potential explanation to the decline in function of this organelle, we consider that recently proposed mechanisms including decreased PINK1 expression, STFPC mutations, or SIRT3 deficiency offer new insights into this finding and should be further study.

Moreover, multiple studies have reported altered proteostasis in aging and IPF lungs, associated with accumulation of aggregated misfolded proteins, defective autophagy, ER stress, and declined of proteasome activity (14, 20, 38). Taken that into consideration, we first used a new molecular rotor detection system to show the intracellular accumulation of misfolded proteins in chronically injured MLE12 cells. Further, we demonstrated that chymotrypsin-like, trypsin-like and caspase-like activities were significantly reduced in these cells, suggesting that impaired proteasome function might contribute to the accumulation of aggregated proteins in our *in vitro* model. We hypothesized that the decline reported in proteasome activity is related to multiple factors previously reported, such as redox alterations by the increase in ROS production, misfolded proteins with toxic oligomers, impaired activation of Nrf1 (a major regulator of proteasome subunits expression) or compromised nuclear translocation of this transcription factor. Furthermore, we previously proposed that this decline in proteasomal activity is detrimental for cellular homeostasis and leads to loss of mitochondria quality control in age-related lung fibrosis by affecting mitochondrial biogenesis, fission/fusion, and mitophagy. Therefore, targeting the UPS offers an attractive therapeutic approach to restore proteostasis and mitochondrial function in the alveolar epithelium.

Heat shock proteins (HSPs) are considered cytoprotective elements by participating in conjunction with the proteasome to degrade misfolded/aggregated proteins. Previous studies have reported reduced Hsp70 levels in lung fibroblasts of IPF patients (43). Consistent with this, we recently demonstrated that the decline of HSF1 activation plays a central role in reducing this chaperone levels in these cells (10). Additionally, Hsp70-knockout mice demonstrated accelerated pulmonary fibrosis after bleomycin administration (43). In contrast, HSP90 is overexpressed in lung fibroblast of IPF lungs, and the inhibition of this chaperone attenuated the progression of pulmonary fibrosis (46). These studies highlight the potential role of HSPs in the pathogenesis of IPF. However, the role of other mitochondria-located HSPs has not been studied in senescent AE2 cells. Our study demonstrated reduced expression of the mitochondria-chaperones Hsp60, Hsp10, and Hsp75 in chronically injured AE2 cells. Interestingly, given senescence cells have shown altered mitochondrial proteins, our results offer a novel mechanism connecting the decline in mitochondrial function and the presence of dysmorphic mitochondria with the decreased expression of these HSPs in senescence AE2 cells. Additionally, we showed that HSF1 activation, the major regulator of HSPs expression, was significantly reduced in these cells. Reduced expression of HSF1 has been shown to cause senescence by eliciting a proteostasis collapse, thereby compromising the ability of cells to perform essential biological activities. However, it is now appreciated that increased post-transcriptional modification, induced via phosphorylation can also contribute to impaired HSF1 activity (11, 12). In support of this idea, we detected a marked increase in Ser307 phosphorylation in chronically injured AE2 cells, thereby resulting in decline HSF1 transcriptional activation and significant accumulation of aggregated/misfolded proteins throughout the entire cytoplasm, including the perinuclear regions.

A comprehensive understanding of stress response after bleomycin exposure is crucial for developing a robust *in vitro* model. We present here an in-depth investigation of the chronic effect of bleomycin on the transcriptome, metabolome, and lipidome of murine lung epithelial cells (MLE12). We envisage that our investigation of the chronic effect of bleomycin would be particularly useful for the establishment of high-throughput assays using the MLE12 cell line, which has been found to be an excellent model to predict the results of acute toxicity and to study mouse transcriptomic. MLE12 cells are one of the best-characterized lung cell lines, and the most recent application of transcriptomic methods on MLE12 cells reported coverage of 5593 DEGs. They are thus a reliable system to obtain information on networks of pathways affected by pro-fibrotic factors that could lead to pulmonary fibrosis.

Thus, we performed RNA sequencing analysis in controls and chronically injured MLE12 cells. A number of DEGs (>8,484) were discovered between controls and chronically injured based on transcriptome datasets. Next, we used a recently published single-cell RNA-sequencing data set to further compared expression in IPF versus healthy lungs. Consistent with previous reports in IPF epithelial cells, our findings identified the activation of transcriptional pathways associated with cell injury and repair in chronically injured AE2 cells. (27, 56). There are similar findings in the transcriptome analysis between our data and one of IPF epithelial cells (32, 56). For instance, cell junction/extracellular matrix organization, response to wounding, p53, PI3K/AKT, and WNT pathways were upregulated, while lipid synthesis, ubiquitin-mediated proteolysis and endosomal protein processing were downregulated. This is all likelihood not coincidental as the exposure to bleomycin has been linked to an increased DNA damage response (34, 37). Other pathways altered in our study correlate with other well-known effects of bleomycin. For instance, DNA replication, DNA repair, and histone modification were downregulated. Nonetheless, there was still opposite results in HIPPO/YAP pathway, and more different in other aspects. We recognize that one limitation of this study is that the changes induced by bleomycin were measured at a late time point and may not reflect the first wave of transcriptional changes induced by this chemical.

Our study also provides mechanistic insights into metabolic alterations induced by a very low concentration of bleomycin and showed putative biomarkers, which are known to be present in IPF lungs. The expected hallmarks of metabolites alterations in the IPF lung were detected in the molecular profiles of MLE12 cells exposed to bleomycin. These changes include upregulation in amino acids (phenylalanine, proline and threonine), and pyrimidine metabolism (uracil), and downregulation in glucose and succinic acid. (18, 59). Moreover, in this study, we provide evidence that exposure to chronic bleomycin altered lipid metabolism in murine AE2 cells, which correlated with findings obtained from metabolomics analysis in IPF lungs (57). Notably, this study confirms increased glycerophospholipids, p-glycerol, and sphingolipids in response to chronic bleomycin. It is not surprising that we found alterations in the lipid metabolism pathway since downregulation in these pathways has been shown in response to bleomycin-induced lung fibrosis. The biological significance of these changes is not clear and requires further exploration.

Overall, the transcriptome and metabolome signatures that we have identified in response to low concentration of bleomycin in ML12 cells are similar to what occurs in IPF lungs. Therefore, our model can help to understand the molecular signature associated with the pathogenesis of lung fibrosis.

---

**Table Supplementary 1:**
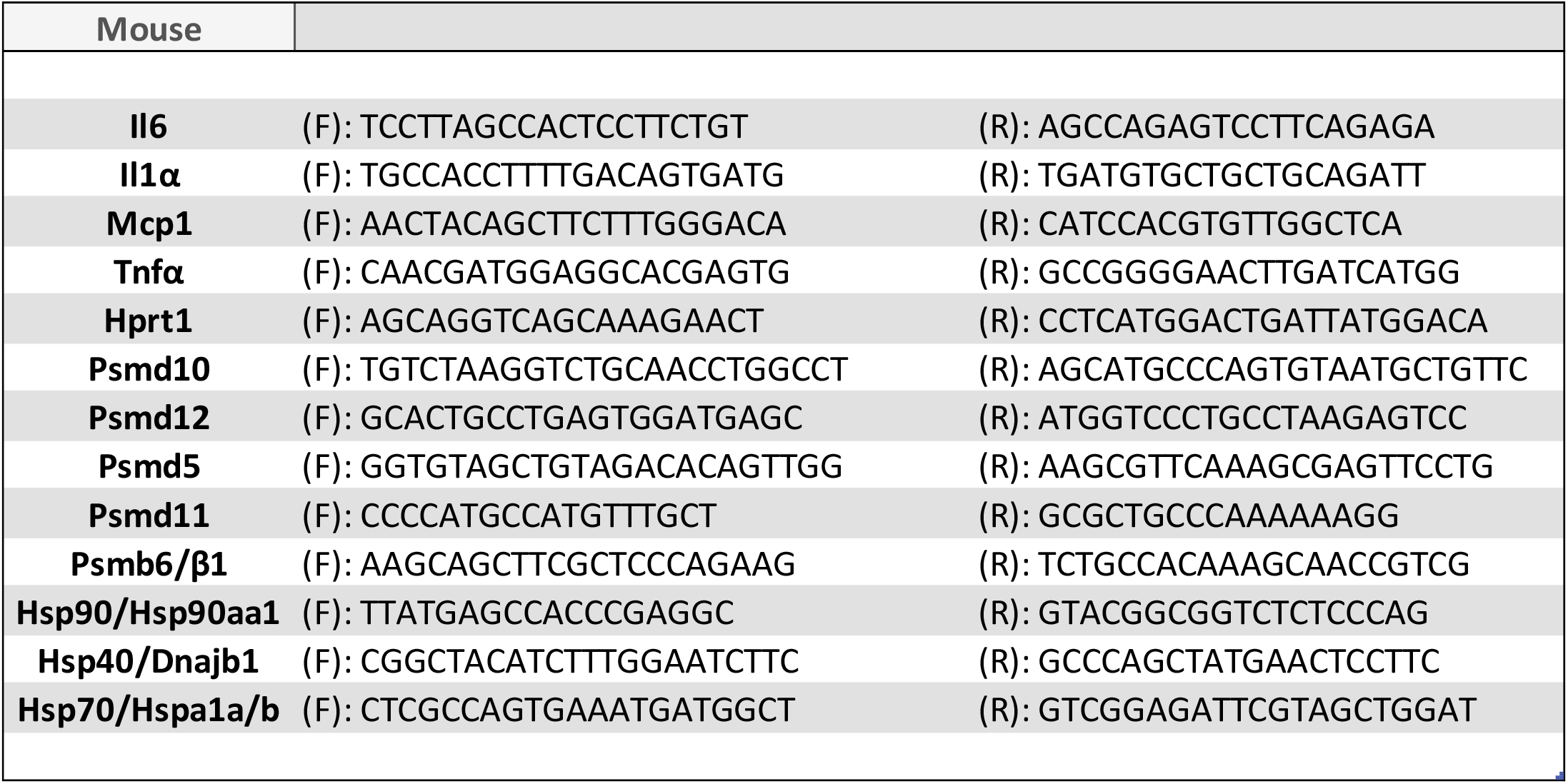
Primers used for qRT-PCR.

**Table Supplementary 2:** The top 20 enriched GO terms in upregulated and downregulated genes from control to the treated/IPF condition.

**A**
| ID | Description | p.adjust |
| --- | --- | --- |
| GO:0070268 | cornification | 7.057E-05 |
| GO:0008544 | epidermis development | 7.057E-05 |
| GO:0031424 | keratinization | 1.595E-04 |
| GO:0030216 | keratinocyte differentiation | 4.138E-04 |
| GO:0043588 | skin development | 4.138E-04 |
| GO:0031581 | hemidesmosome assembly | 4.506E-04 |
| GO:0034330 | cell junction organization | 9.310E-04 |
| GO:0034329 | cell junction assembly | 1.109E-03 |
| GO:0007044 | cell-substrate junction assembly | 1.109E-03 |
| GO:0009913 | epidermal cell differentiation | 1.109E-03 |
| GO:0043062 | extracellular structure organization | 3.662E-03 |
| GO:0030198 | extracellular matrix organization | 4.869E-03 |
| GO:0048660 | regulation of smooth muscle cell proliferation | 5.397E-03 |
| GO:0048659 | smooth muscle cell proliferation | 5.397E-03 |
| GO:1990478 | response to ultrasound | 7.811E-03 |
| GO:0001655 | urogenital system development | 1.007E-02 |
| GO:0072001 | renal system development | 2.039E-02 |
| GO:0030574 | collagen catabolic process | 2.308E-02 |
| GO:0048145 | regulation of fibroblast proliferation | 2.308E-02 |
| GO:0030336 | negative regulation of cell migration | 2.308E-02 |

**B**
| ID | Description | p.adjust |
| --- | --- | --- |
| GO:0006695 | cholesterol biosynthetic process | 7.09E-12 |
| GO:1902653 | secondary alcohol biosynthetic process | 7.09E-12 |
| GO:0016126 | sterol biosynthetic process | 1.10E-11 |
| GO:0045540 | regulation of cholesterol biosynthetic process | 2.08E-09 |
| GO:0106118 | regulation of sterol biosynthetic process | 2.08E-09 |
| GO:0008203 | cholesterol metabolic process | 2.08E-09 |
| GO:1902652 | secondary alcohol metabolic process | 2.33E-09 |
| GO:0050810 | regulation of steroid biosynthetic process | 2.33E-09 |
| GO:0016125 | sterol metabolic process | 3.96E-09 |
| GO:0090181 | regulation of cholesterol metabolic process | 1.86E-08 |
| GO:0006694 | steroid biosynthetic process | 2.41E-08 |
| GO:0046890 | regulation of lipid biosynthetic process | 2.99E-08 |
| GO:0019218 | regulation of steroid metabolic process | 3.27E-08 |
| GO:0062012 | regulation of small molecule metabolic process | 4.72E-08 |
| GO:0008202 | steroid metabolic process | 7.81E-08 |
| GO:0046165 | alcohol biosynthetic process | 9.43E-08 |
| GO:1902930 | regulation of alcohol biosynthetic process | 1.53E-07 |
| GO:0019216 | regulation of lipid metabolic process | 4.22E-07 |
| GO:1901617 | organic hydroxy compound biosynthetic process | 1.22E-06 |
| GO:0006066 | alcohol metabolic process | 2.95E-05 |

**Figure Supplementary 1:**
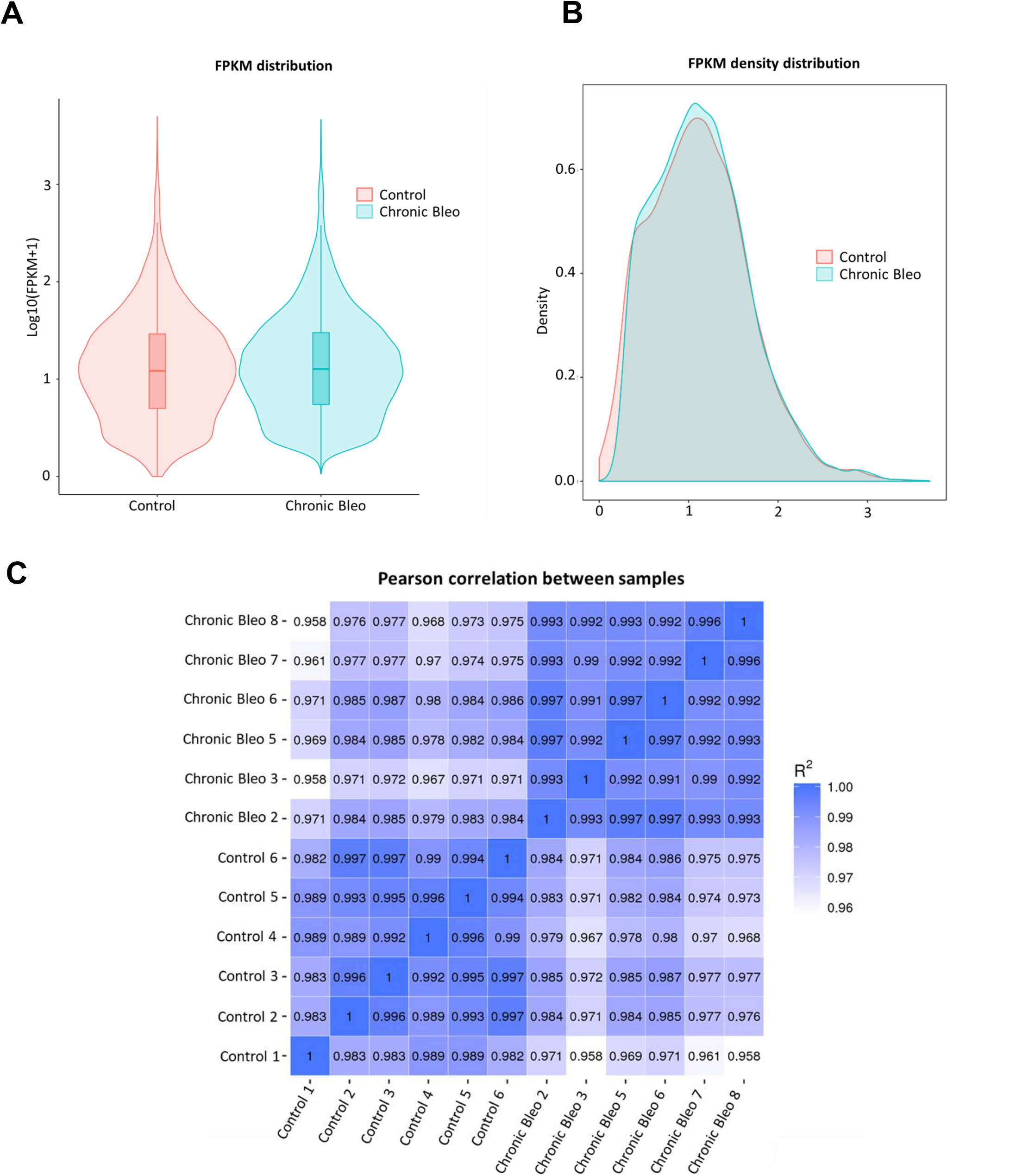
Bioinformatics analysis of RNA-seq data. FRKM distribution of control and chronical injury MLE12 cells. B) FRKM density distribution between the two groups. C) Pearson correlation between samples.

**Figure Supplementary 2:**
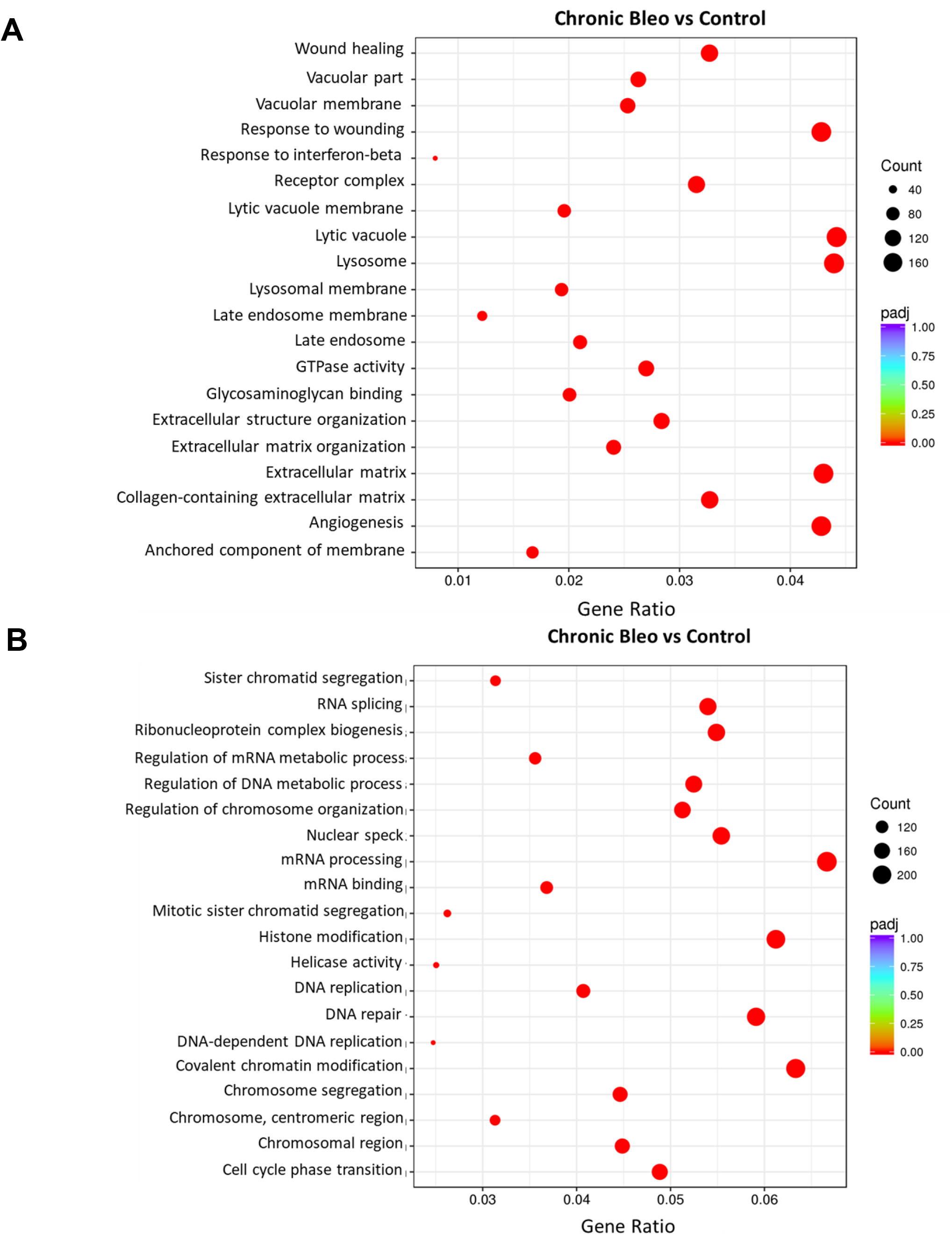
GO pathway analysis of differentially expressed genes. Advance bubble chart shows enrichment of differentially expressed genes in signaling pathways (A) upregulated and B) downregulated.

**Figure Supplementary 3:**
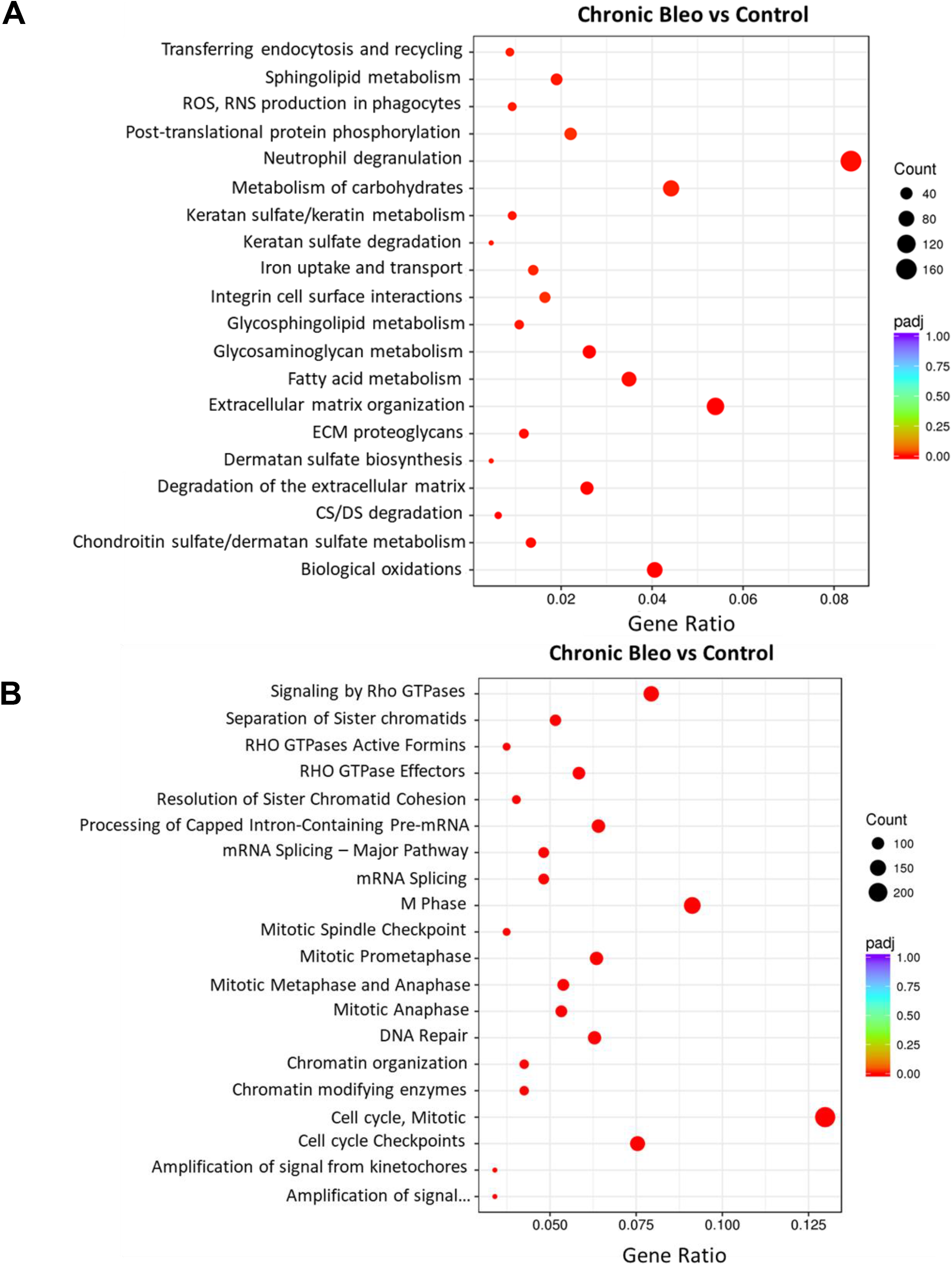
Reactome pathway enrichment analysis of differentially expressed genes. Advance bubble chart shows enrichment of differentially expressed genes in signaling pathways (A) upregulated and B) downregulated.

**Figure Supplementary 4.**
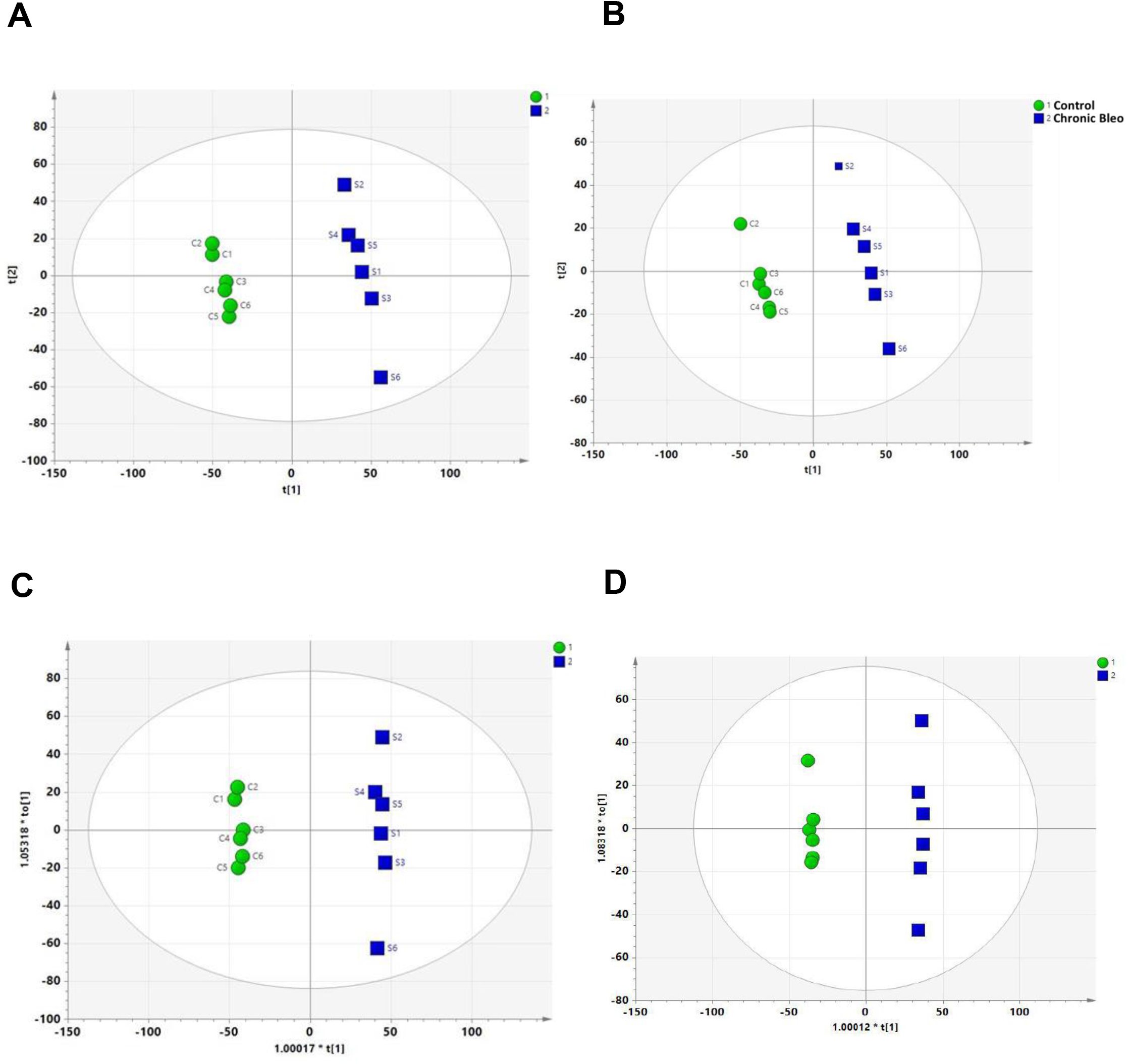
The score plots from PLS-DA and OPLS-DA model. PLS-DA score plot of positive (A) and negative ion mode (B). OPLS-DA score plot of positive (C) and negative ion mode (D).

**Figure Supplementary 5.**
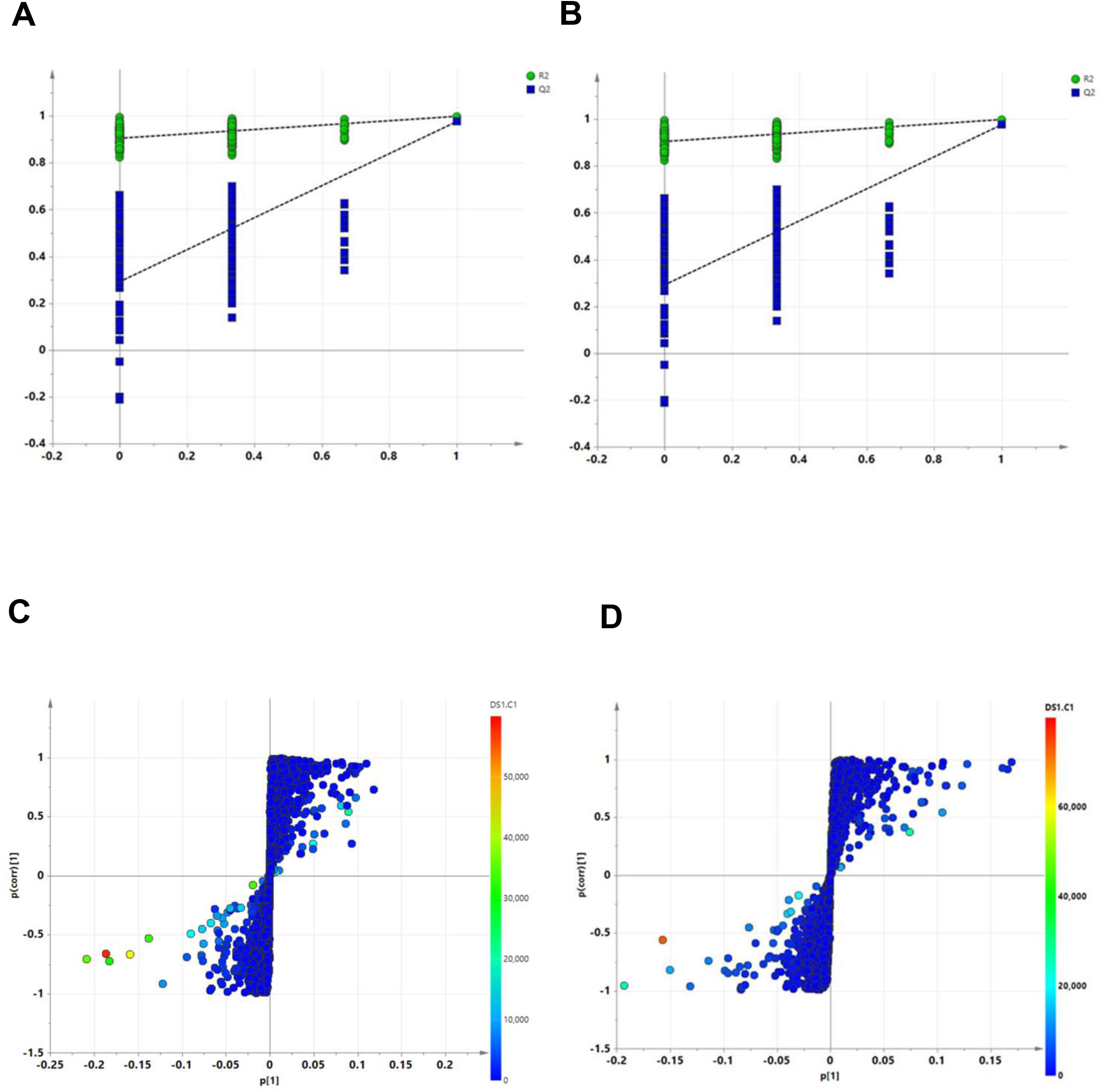
Non-targeted metabolomics analysis from control and chronically injured MLE12 cells in positive and negative mode. Permutation test of PLS-DA model in control (green point) and injured cells (blue point). B) An OPLS-DA loading S-plot. The axis is the modelled co-variation and the y-axis is the loading vector of the predictive component collected from the two groups.

**Figure Supplementary 6.**
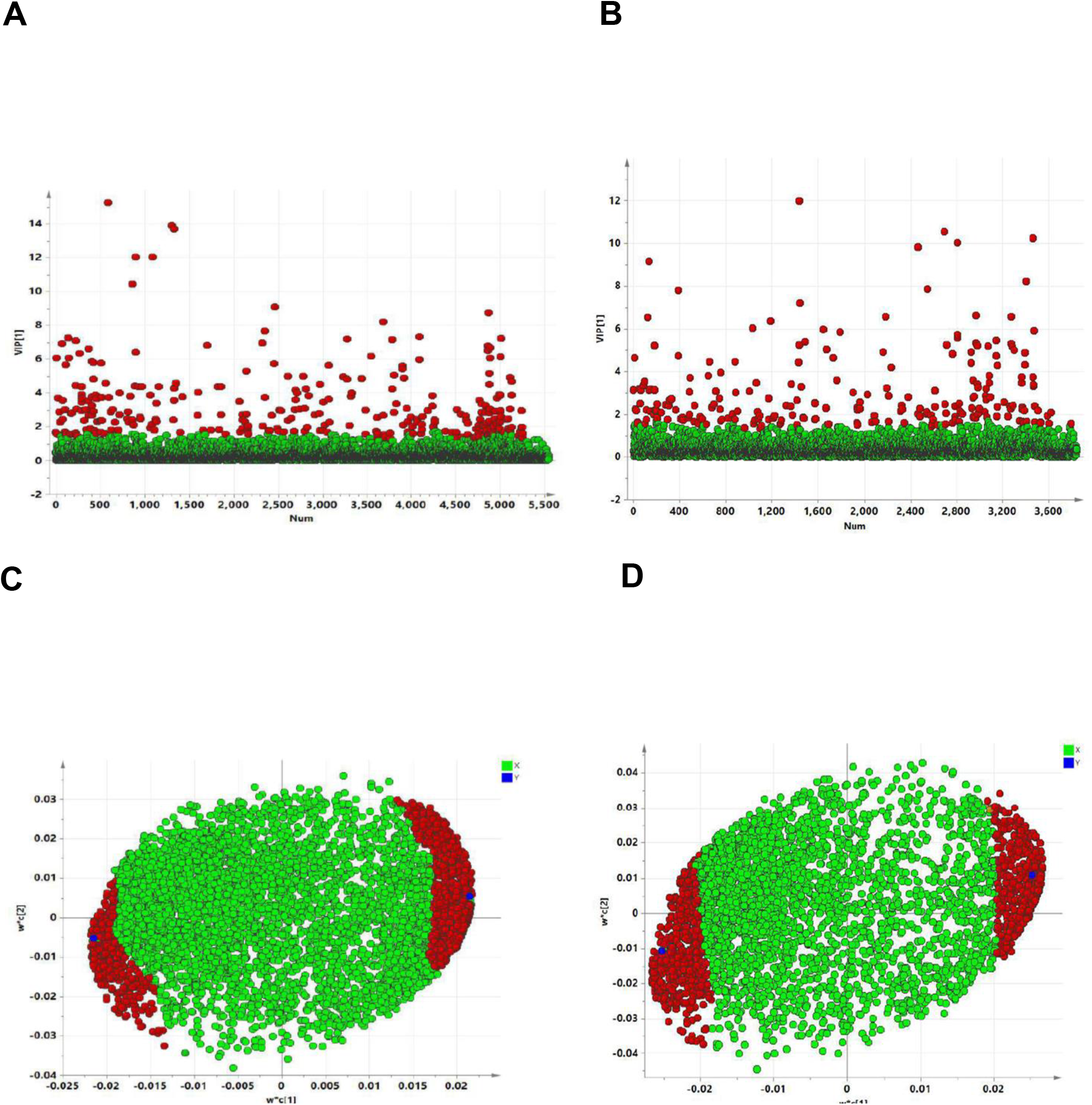
The distribution of VIP values (VIP > 1.5) of the significantly changed metabolites, positive (A) and (B) negative ion mode between the groups. The PLS-DA loading plot, positive (C) and negative ion mode (D) between the control and chronically injured cells. The metabolites with red box were labeled as significant compounds (VIP>1.5) analysis‥

**Figure Supplementary 7.**
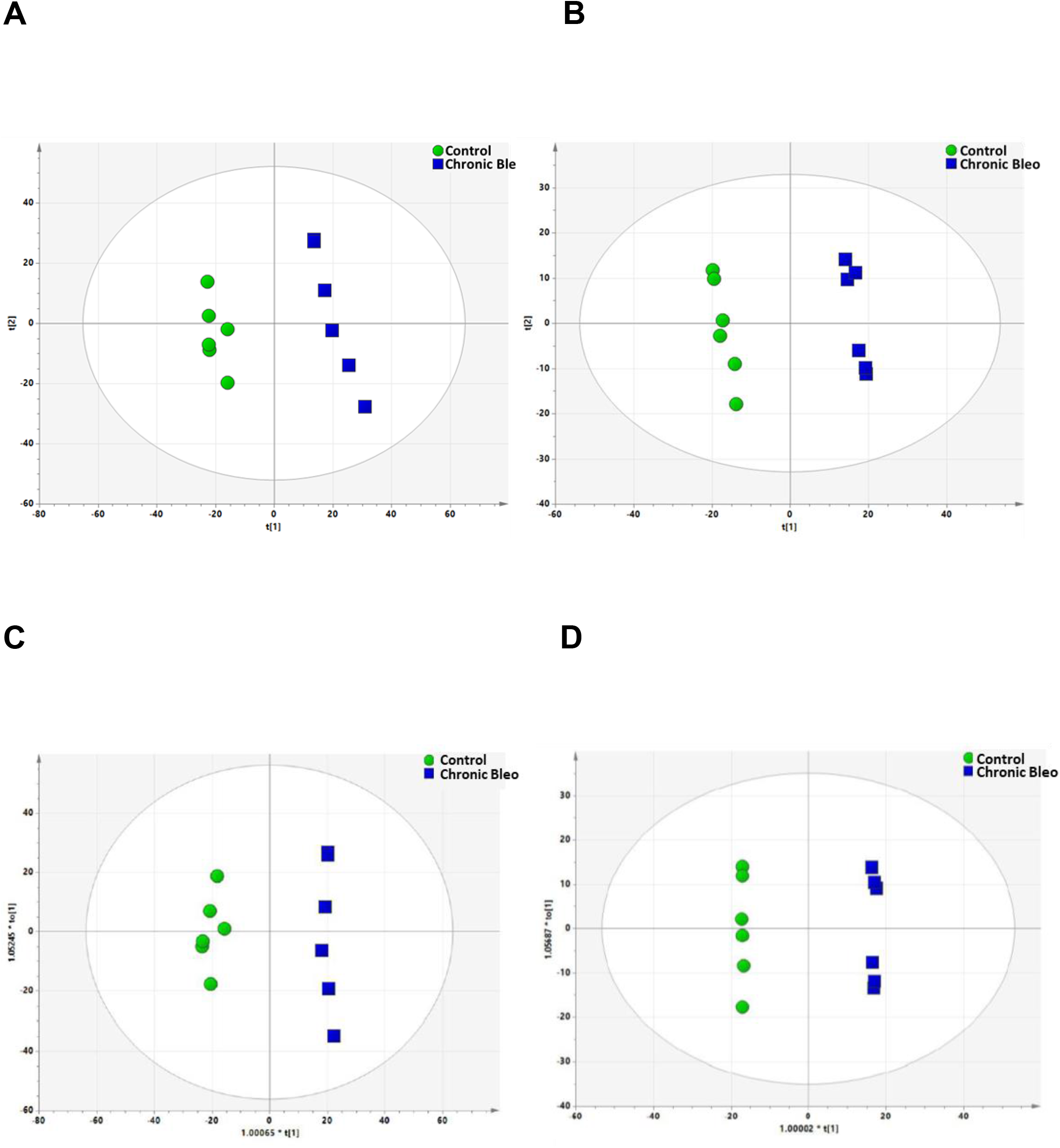
The score plots from PLS-DA and OPLS-DA model, obtained from LC-MS/MS profile of control and bleomycin injured MLE12 cells. PLS-DA score plot of positive (A) and negative ion mode (B). OPLS-DA score plot of positive (C) and negative ion mode (D).

**Figure Supplementary 8.**
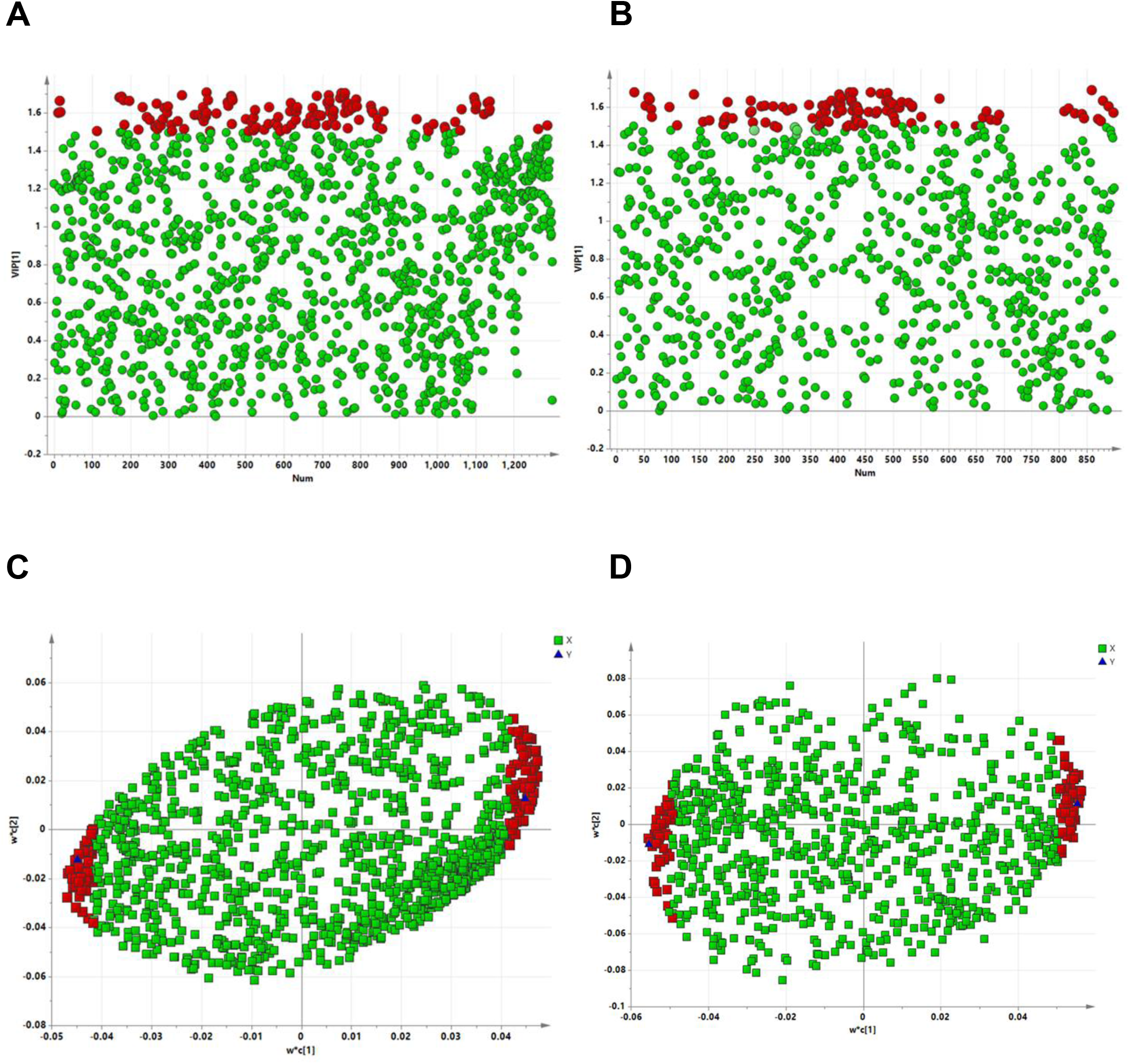
The distribution of VIP values (VIP > 1.5) of the significantly changed metabolites, positive (A) and (B) negative ion mode between controls and chronically injured MLE12 cells. The PLS-DA loading plot, positive (C) and negative ion mode (D) between the control and injured cells. The metabolites with red box were labeled as significant compounds (VIP>1.5) analysis‥

